# Prospective pan-cancer phosphoproteomics at clinical scale extends therapeutic options in precision oncology

**DOI:** 10.64898/2026.07.08.737171

**Authors:** Annika Schneider, Julia Wortmann, Cecilia Bang Jensen, Leonardo Estrada Duenas, Maria-Veronica Teleanu, Amirhossein Sakhteman, Firas Hamood, Florian P. Bayer, Christoph Stange, Jeremiah Bintang Santoso, Jennifer Huellein, Nindo Punturi, Lee Dolat, Peter Horak, Moritz Resch, Nicole Kabella, Stefanie Höfer, Simon Kreutzfeldt, Christoph E. Heilig, Maximilian Werner, Chen Hong, Barbara Hutter, Katja Beck, Eva Reisinger, Katrin Pfütze, Chien-Yun Lee, Yun-Chien Chang, Christel Herold-Mende, Malgorzata Oles, Kathrin Schramm, Stephanie Wilhelm, Andreas Unterberg, Katja Steiger, Carolin Mogler, David Jones, Olaf Witt, Daniel Hübschmann, Ulrich Keilholz, Damian Rieke, Frederick Klauschen, Albrecht Stenzinger, Sebastian Bauer, Jens Thomas Siveke, Christian Brandts, Thomas Kindler, Evelin Schröck, Melanie Boerries, Michael Bitzer, Klaus Schulze-Osthoff, Tobias Herold, Rainer Claus, Roland Ullrich, Florian Lueke, Verena I. Gaidzik, Ralf Bargou, Vivek Venkataramani, Pauline Wimberger, Theresa Link, Verena Kiver, Sabine Heublein, Nina Ditsch, Wolfgang Janni, Verena Thewes, Marc Zapatka, Mario Hlevnjak, Andreas Schneeweiss, Peter Lichter, Daniel M. Freed, Wilko Weichert, Hanno Glimm, Stefan M. Pfister, Stefan Fröhling, Matthew The, Bernhard Kuster

**Author notes:** these authors contributed equally (Annika Schneider, Julia Wortmann, Cecilia Bang Jensen, Leonardo Estrada Duenas, Maria-Veronica Teleanu). shared senior authors (Stefan Fröhling, Matthew The, Bernhard Kuster).

## Abstract

Genomics-guided precision oncology has improved survival in cancer entities with actionable mutations but cannot capture oncogenic signaling that manifests at the protein level. Here, we report a prospective, real-world pan-cancer study profiling proteomes and phosphoproteomes of 1,998 tumor samples from adults and children with rare or advanced cancers enrolled in the German precision oncology programs DKFZ/NCT/DKTK MASTER, CATCH and INFORM and their molecular tumor boards (MTBs). We developed tumor proteome activity status (TOPAS) scores for 46 clinically relevant kinases, an immune activity score capturing antigen presentation and T-cell activation and identified therapeutically targetable cell-surface proteins for 94% of patients. These readouts enhance MTB recommendations by exposing actionable non-genomic kinase activity, refining interpretation of oncogenic genome alterations, and highlighting cell-surface treatment options. Three proof-of-concept analyses indicate clinical utility including kinase activity-stratified pazopanib response in sarcoma, immune activity score-tracked checkpoint-inhibitor outcomes pan-cancer, and a phosphoproteomic biomarker distinguishing EGFR-inhibitor response in chordoma.

## INTRODUCTION

Genomics-based precision oncology has delivered survival benefits where therapeutically actionable alterations are well-defined. Examples include kinase inhibitors in fusion- or mutation-driven cancers, HER2-directed therapies in amplified tumors, and tumor-agnostic approval for NTRK fusions and MSI-H disease across histologies. Yet the scope of actionability remains narrow: across multiple large-scale prospective sequencing programs, only 20–30% of patients harbor alterations with established therapeutic relevance, and objective response rates in matched cohorts rarely exceed 15–20%^1-5^. Similarly, in the MTBs of the MASTER^6,7^ and INFORM^8,9^ programs, benefit from molecularly matched treatment was confined to a minority of patients with highly actionable genomic findings, while patients without those only occasionally benefited. This partly reflects context-dependent differences in tumorigenesis, including tissue of origin, tumor microenvironment, and co-occurring genomic alterations, as well as the fact that oncogenic signaling driven by non-genomic mechanisms may be equally relevant for treatment selection^10^. The diagnostic challenge is exacerbated in advanced or heavily pretreated patients, where long tumor evolutionary trajectories make assessing actionability from genomes alone particularly difficult^11^.

Kinase dysregulation is a defining hallmark of cancer, and kinase inhibitors (KIs) rank among the most successful targeted therapies today^12^. Still, current precision oncology protocols rely almost exclusively on genome and, rarely, transcriptome profiling, and their interpretation is typically limited to known or presumed oncogenic alterations. For kinases, these include genomic amplifications, gain-of-function mutations, and chromosomal rearrangements, all of which are thought to precipitate oncogenic downstream effects. However, kinases are regulated beyond gene expression such as phosphorylation of functionally important amino acids, co-factor availability, scaffold interactions, and substrate competition. Genomic surrogates for aberrant kinase activity, therefore, carry inherent limitations, particularly in complex tumor tissues^13–15^. The presence of a kinase-activating alteration neither necessarily produces hyperactive kinase signaling nor does its absence rule it out. Furthermore, most genomic alterations are functionally not understood, and even those that are, can have variable effects depending on cellular context^16,17^. Additional layers of regulation relevant to oncogenesis, including epigenetic changes and post-translational modifications, also go undetected by standard genome or transcriptome profiling. Collectively, this often makes it impossible to predict the consequence of nucleotide-level alterations in a particular patient, underscoring an urgent need for diagnostics that measure oncogenic signaling directly at the protein level.

Proteome and phosphoproteome profiling (from here collectively referred to as phosphoproteome profiles) of clinical tumor samples has emerged as a complementary molecular layer to address these limitations. The Clinical Proteomic Tumor Analysis Consortium (CPTAC) has produced foundational resources across more than ten cancer types, demonstrating that integrating phosphoproteomics with genomic data uncovers druggable targets and kinase-driven biology not apparent from genome or transcriptome profiling alone^18–20^. Computational frameworks including KSEA, INKA and KSTAR have established that phosphorylation site (p-site) profiles can be condensed into kinase activity scores informative of signaling states, outperforming genomic surrogates for predicting kinase inhibitor response in cell lines and patient-derived models^21–26^. However, existing resources were almost exclusively derived from retrospectively collected, treatment-naive surgical resections from common cancer types, and data interpretation pipelines were optimized for cohort-level rather than individual-patient interpretation. Here, we demonstrate that prospective phosphoproteome profiling is feasible at clinical scale and can be integrated into routine MTB workflows. Applied to rare and heavily pretreated cancers, where unmet clinical need is greatest, we show that phosphoproteomics yields clinically actionable insights not accessible from genomic data alone.

## RESULTS

### Integration of phosphoproteome profiling into prospective MTB workflows is feasible at clinical scale

To augment clinical decision-making in MTBs with phosphoproteome-wide profiles, we established a laboratory workflow that enables a two-week turnaround from sample receipt to MTB discussion (**Figure 1A**, Methods). Briefly, the proteome was extracted from tumor tissue, digested into peptides, and samples were multiplexed into batches by stable isotope labeling (tandem mass tags, TMT) Each batch was fractionated by high-pH reversed-phase chromatography (bRPLC), phosphopeptides (p-peptides) were enriched by immobilized metal affinity chromatography (IMAC), and proteomes and phosphoproteomes were analyzed by liquid chromatography–tandem mass spectrometry (LC-MS/MS). A custom-developed automated data processing pipeline performed missing-value reduction, quality filtering, normalization, annotation, and activity scoring (accompanying manuscript: TOPAS: phosphoproteome data analysis and decision support platform for molecular tumor boards). The pipeline has been executed weekly since June 2022, building a continuously growing background cohort so that each new case benefits from, and contributes to, an ever-expanding reference (**Figure 1B**). Two internal reference samples included in each batch confirmed that proteome and phosphoproteome quantification remained fit-for-purpose over a five-year period (**Figures S1A and B**).

**Figure 1:**
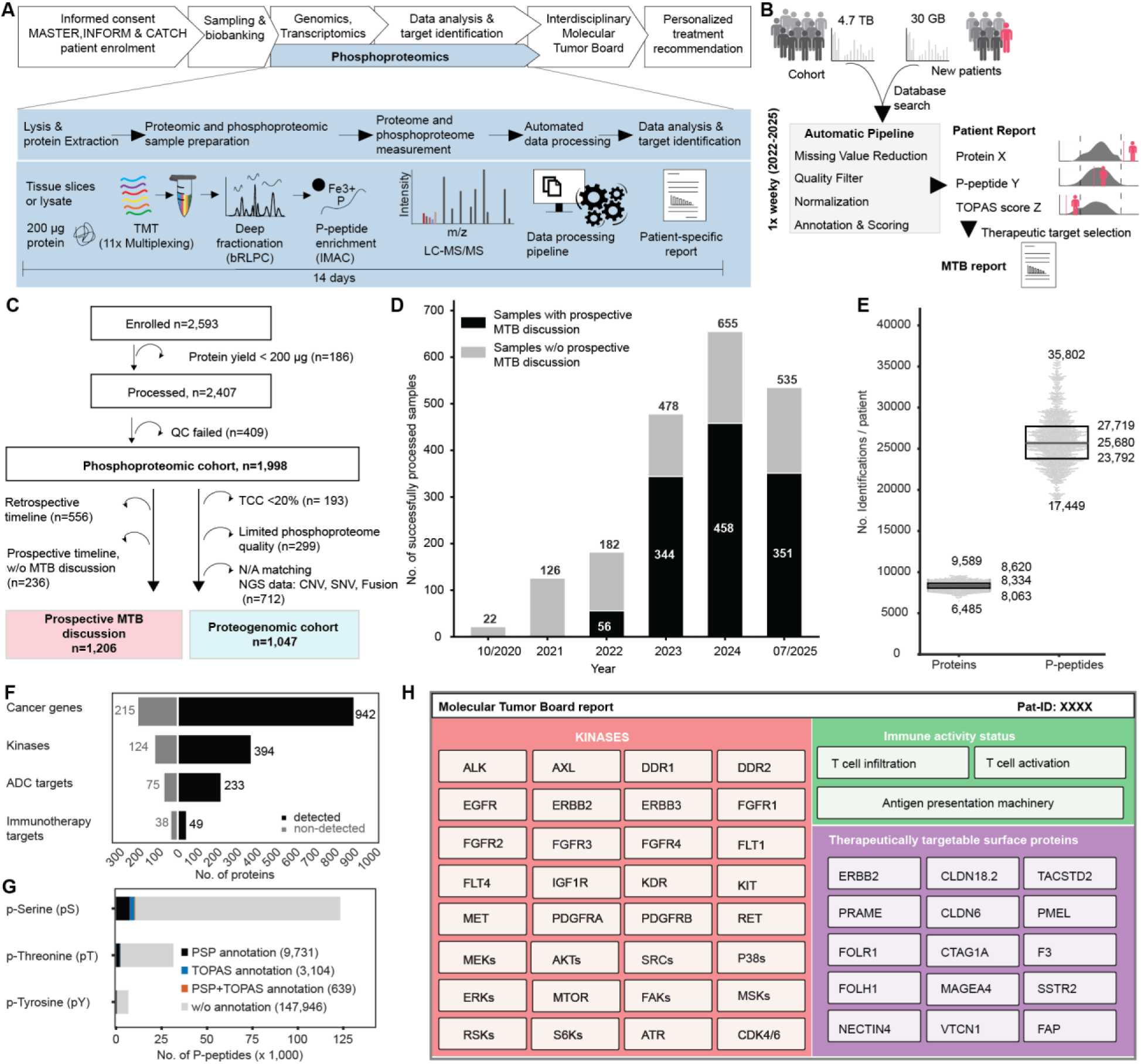
(Phospho)proteome profiling in precision oncology trials. (A) Workflow and timing of phosphoproteome profiling integrated into the clinical workflows of the MASTER, INFORM, and CATCH precision oncology trials. (B) Schematic of the in-house data processing pipeline that automatically generates molecular tumor board (MTB) reports based on a weekly growing pan-cancer cohort. (C) Consort flow diagram depicting the progression of tissue samples through the phosphoproteome profiling workflow, from sample shipment to MTB discussion or inclusion in proteogenomic analyses. (D) Number of QC-passing tumor samples per year with (black) or without (grey) prospective MTB discussion. (E) Distribution of the number of proteins and p-peptides (identified at 1% FDR) quantified per tumor sample. (F) Number of detected (black) versus undetected (grey) proteins across four protein-of-interest groups: OncoKB cancer genes, human kinases, ADC targets, and immunotherapy targets. (G) Distribution of phosphoserine (pS), phosphothreonine (pT), and phosphotyrosine (pY) residues, with or without tumor proteome activity status (TOPAS) and/or PhosphoSitePlus (PSP) annotation, across all detected p-peptides. (H) Schematic MTB report overview depicting reported biomarkers for three drug categories: kinase inhibitors, immune checkpoint inhibitors, and surface expression-targeting drugs.

Between October 2020 and July 2025, 2,593 tumor tissue samples from numerous centers across Germany participating in the DKFZ/NCT/DKTK MASTER or CATCH trials, and the pediatric precision oncology INFORM registry were collected in two centralized and accredited sample processing laboratories. 2,407 samples yielded sufficient protein (≥200 µg) and 1,998 passed quality-control (QC) criteria (**Figure S1C**). Overall, 72% (n=1,442) of samples met a prospective timeline and 84% of those (n=1,206) were discussed prospectively in MTBs. For proteogenomic analyses, 1,047 samples with tumor cell content ≥20% (**Figures S1D and E**), high phosphoproteome quality (**Figure S1F**), and matched whole-genome or exome sequencing data were included (**Figure 1C, Table S1**). Over time, sample accrual increased steadily as did the number of weekly cases discussed in MTBs, reflecting increasing workflow automation and maturation of MTB reporting (**Figure 1D**).

Between 6,485 and 9,589 proteins were quantified per patient (median 8,334; interquartile range, IQR = 8,063–8,620; total: 13,069, **Figure 1E**). The phosphoproteome covered 17,449 to 35,802 phospho-peptides (p-peptides) per patient (median 25,680; IQR = 23,792–27,719; total: 148,224, **Figure 1E**). The collective data showed broad coverage of proteins of direct relevance to cancer biology, notably OncoKB-annotated cancer genes (942/1,157), protein kinases (394/518), antibody–drug conjugate (ADC) targets (233/308 approved, in trials, or in development), and immunotherapy targets (49/87 approved, in trials, or in development; **Figure 1F**). Ubiquitously expressed proteins (referred to as low specificity proteins according to the tissue specificity classification in the human proteome atlas^27^ [HPA], such as MAPK1 and PTEN) were detected in virtually every patient, whereas tissue-enriched proteins, such as MAGEA4 and MS4A1, were detected in a subset of patient samples only (**Figure S1G**).

Among all detected p-sites, 76.4% were located on serine (pS), 19.4% on threonine (pT), and 4.2% on tyrosine (pY) residues. Only 6.4% had a functional annotation in PhosphoSitePlus (PSP) and just 2.5% were annotated with one or more upstream kinases using DecryptM data^28^ (**Figure 1G**). The large number, but modest overlap, of p-peptides between patients is consistent with the substantial molecular heterogeneity of the underlying tumor tissues (**Figure S1H**). Still, the phosphoproteome captured many clinically relevant and actionable signaling nodes across multiple druggable axes: Receptor tyrosine kinases (RTKs), key effectors of the MAPK and PI3K–AKT–mTOR cascades, as well as cell-cycle regulators, DNA-damage response kinases, stress kinases and non-receptor tyrosine kinases (**Fig. 1H, Table S2**). Beyond kinase signaling, the dataset captured tumor-reactive T-cell biology including T-cell infiltration (e.g. CD3D/E/G, CD8a), T-cell activation (e.g. CD8a S231, PDCD1 S267, STAT1 Y701, CD38), and the antigen-presentation machinery (e.g. TAP1, TAPBP, PSMB9, CDC1c), as well as therapeutically targetable cell-surface proteins such as ERBB2, PRAME, FOLR1, CTAG1A (NY-ESO-1), CLDN18.2, TACSTD2 (Trop-2) and NECTIN4, among others (**Table S2**). Together, these readouts were used to populate patient-specific reports generated for each MTB meeting (**Figure 1H**). To ensure clinical-grade reliability, every reported value was gated by standardized QC-criteria, including variance ratio thresholds, minimum peptide counts, and TMT intensity bounds, with per-patient QC summaries provided alongside each report (**Figure S1I, S1K and S1J**, **Methods**).

### Tissue-of-origin dominates protein expression levels and enables robust subtype classification

The cohort comprises 1,998 samples representing 194 unique cancer subtypes (by oncotree-code^29^) across 32 tissues of origin, broadly classified into sarcoma (n=965), carcinoma (n=756), neuroendocrine neoplasms (NET/NEC, n=93), and other entities (n=184) including CNS tumors, melanoma, germ cell tumors and thymoma (**Figures 2A and S2A**). This extreme histological diversity reflects the clinical focus of the DKFZ/NCT/DKTK MASTER, INFORM, and CATCH programs. Theyspan the full breadth of oncology practice: pediatric (INFORM) and adult patients, ranging from ultra-rare malignancies (MASTER) to common solid tumors in advanced or refractory stages (CATCH). Across all three programs, standard treatment options were frequently exhausted and functional molecular insights were most urgently needed. Accordingly, most samples (72%) were derived from metastatic lesions across diverse tissue topologies including liver, soft tissue, lung, bone, and lymph node, and 74% met the European Union definition of rare cancers (<6 per 1,000,000; **Figure 2B**). The broad distribution of tumor cell content (0-100%) further underscores the real-world clinical reality under which these samples were collected. Unsupervised dimensionality reduction (UMAP) of patient samples revealed that proteome and phosphoproteome abundance levels mostly describe cancer subtypes (**Figure 2C**). We identified optimized proteome signatures describing cancer subtypes (**Figure S2B**), major histology classes (**Figure S2C**), and less dominant sample features like metastatic site (**Figure S2D**), indicating that tissue-of-origin proteomic signatures persist across metastatic sites, variable tumor purity, and diverse clinical histories.

**Figure 2:**
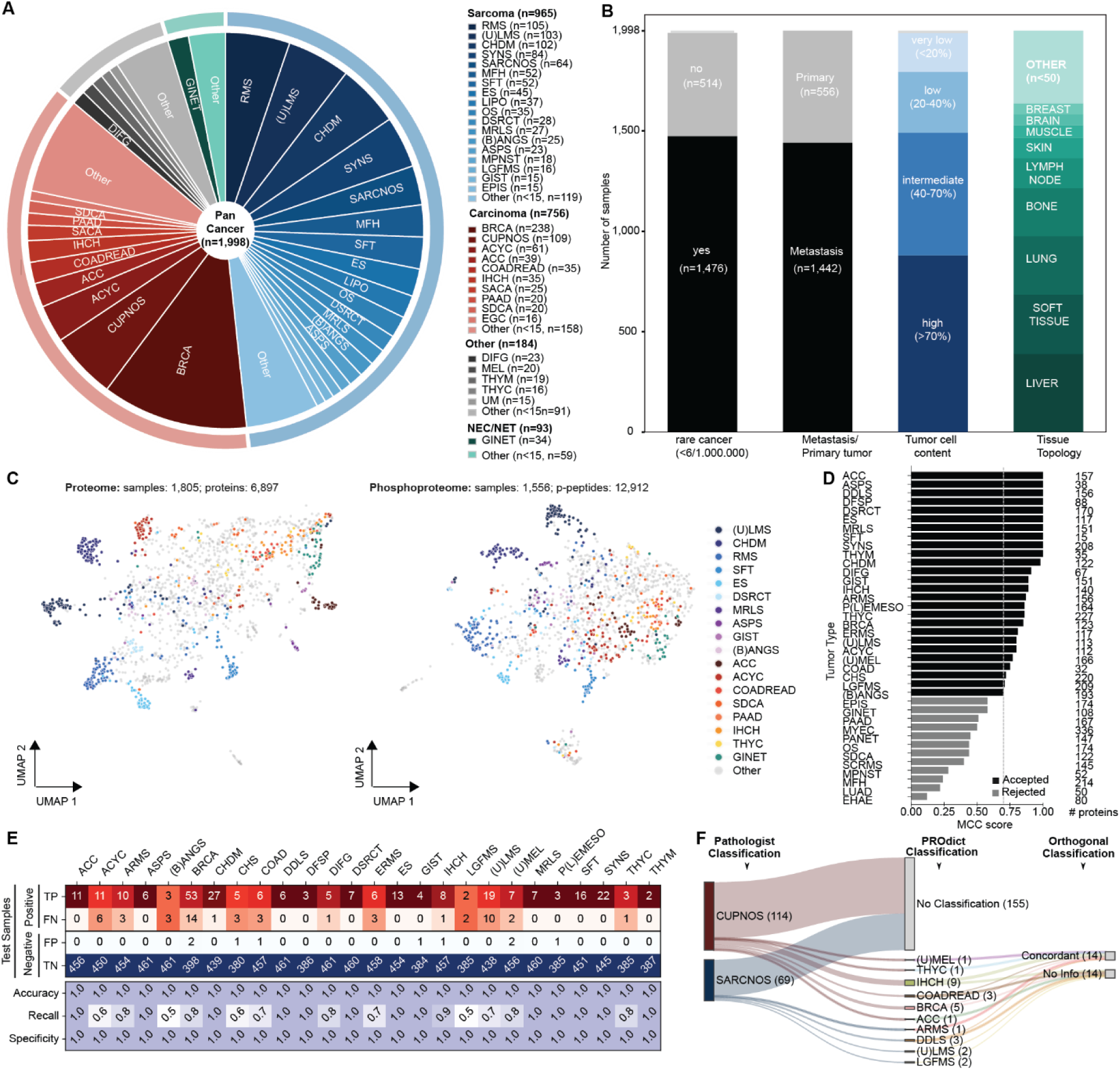
Proteome-based robust cancer subtype classification of a real-world pan-cancer cohort. (A) Pie chart depicting the composition of the pan-cancer cohort by cancer subtype (OncoTree-based classification) across carcinoma (red), sarcoma (blue), neuroendocrine neoplasms (green), and others (grey). (B) Stacked bar graphs depicting the fractions of rare cancer specimens, metastatic specimens, tumor purity categories, and tissue topologies. (C) UMAP projection of expression data from 6,897 proteins (>80% occurrence; left) and 12,912 p-peptide groups (>80% occurrence; right), colored by cancer subtype. (D) Bar chart depicting the Matthews Correlation Coefficient (MCC) score for all 38 trained models (one per cancer subtype with ≥10 samples). Retained models are shown as black bars and rejected models as grey bars, the vertical dashed line indicates the threshold for acceptance (MCC > 0.7). The number of proteins used per model is indicated on the right side of the chart. (E) Classification confusion matrix for all 26 retained models on test samples, accompanied with classification evaluation metrics such as Accuracy, Recall and Specificity for each model. (F) Sankey diagram depicting the re-classification of CUPNOS (red) and SARCNOS (blue) tumor specimens using the 26 available models (middle). Each re-classification was evaluated for orthogonal support using other available omics data (right). *RMS = Rhabdomyosarcoma, (U)LMS = (Uterine) Leiomyosarcoma, CHDM = Chordoma, SYNS = Synovial sarcoma, SARCNOS = Sarcoma, not otherwise specified (NOS), MFH = Undifferentiated Pleomorphic Sarcoma/Malignant Fibrous Histiocytoma/High-Grade Spindle Cell Sarcoma, SFT = Solitary Fibrous Tumor, ES = Ewing Sarcoma, LIPO = Liposarcoma, OS = Osteosarcoma, DSRCT = Desmoplastic Small Round Cell Tumor, MRLS = Myxoid/Round-Cell Liposarcoma, (B)ANGS = (Breast) Angiosarcoma, ASPS = Alveolar Soft Part Sarcoma, MPNST = Malignant Peripheral Nerve Sheath Tumor, LGFMS = Low-Grade Fibromyxoid Sarcoma, GIST = Gastrointestinal Stromal Tumor, EPIS = Epithelioid Sarcoma, BRCA = Breast Carcinoma, CUPNOS = Cancer of Unknown Primary, NOS, ACYC = Adenoid Cystic Carcinoma, ACC = Adrenocortical Carcinoma, COADREAD = Colorectal Adenocarcinoma, IHCH = Intrahepatic Cholangiocarcinoma, SACA = Salivary Carcinoma, PAAD = Pancreatic Adenocarcinoma, SDCA = Salivary Duct Carcinoma, EGC = Esophagogastric Adenocarcinoma, DIFG = Diffuse Glioma, MEL = Melanoma, THYM = Thymoma, THYC = Thymic Carcinoma, UM = Uveal Melanoma, GINET = Gastrointestinal Neuroendocrine Tumor, NEC = Neuroendocrine Carcinoma, NET = Neuroendocrine Tumor*.

We exploited this diversity to build a proteome-based cancer subtype classifier. Prior machine learning models^30,31^ were mostly trained on non-metastatic tumors and often excluded rare subtypes, limiting their utility in the clinical scenarios where they are most needed^32–34^. Instead, we developed PROdict, a proteome-based classifier trained on 1,155 tumor samples spanning 180 subtypes and 39 tissue sites, with 68% of samples derived from metastatic lesions (**Figure S2E**). For each tumor subtype with ≥10 samples, a one-vs-all binary classifier was constructed. Proteins with high discriminative power were preselected using logistic regression with elastic net regularization, followed by final model training with ridge-regularized logistic regression (**Figure S2F).** Among the **38** cancer subtype models evaluated, **26** exhibited robust predictive performance (Matthews Correlation Coefficient, MCC >0.7) in an independent validation cohort (n = 467, 104 subtypes; **Figure 2D**) and were retained for downstream analysis. Successful classifiers relied on compact, interpretable protein signatures ranging from 15 to 227 proteins per subtype (**Figure S2G**). In combination with a precision-focused probability threshold for clinically robust reporting (**Figure S2H**), PROdict achieved correct classification of 265/365 samples, while minimizing incorrect assignment to other subtypes (9 false positives, **Figure 2E**). Interestingly, PROdict suggested a specific cancer subtype for 8 of 77 SARCNOS cases and 20 of 134 CUPNOS cases (sarcoma/cancer not otherwise specified; **Figure 2F**), underscoring its utility for resolving diagnostically challenging tumors. In 14 of 28 (50%) reclassified cases, concordant evidence for the re-diagnosis was provided by other omics layers (genomics/transcriptomics/methylomics) or clinical history (**Figure 2F, Table S3**). These results establish PROdict as the first proteome-based classifier to robustly identify 26 clinically relevant cancer subtypes in a real-world, metastatic pan-cancer cohort. While the diversity of tumor biology captured here is substantial, the breadth of cancer subtype diversity in clinical practice far exceeds current coverage, highlighting the need for continued expansion.

### Pan-cancer surface proteome profiling expands the actionable target space in rare and advanced cancers

Therapies that engage cell-surface proteins have entered a massive expansion phase, with 19 antibody therapeutics gaining first approval in 2025 alone, >200 ADC and 1,580 CAR-T ongoing clinical trials, and a commercial late-stage pipeline of >200 antibodies targeting an ever-expanding set of surface antigens^35–37^.The bottleneck for the field has shifted from molecular engineering to a more fundamental biological question: Which surface proteins, in which tumors, at what abundance level, can be productively targeted? Because each therapeutic modality has a distinct expression threshold below which it loses clinical efficacy, target nomination cannot be reduced to a binary present/absent call but requires quantitative, per-tumor abundance measurement. Immunohistochemistry neither scales to the required number of antigens nor returns robust quantitative data, while mass spectrometry-based proteomics achieves both. To map the surface proteome of our cohort, we integrated three independent surface-protein catalogues: the mass-spectrometry-derived Cell Surface Protein Atlas (CSPA)^38^, the antibody-based Human Protein Atlas (HPA)^27^, and the in-silico tool SURFY^39^. Together, these resources list 4,416 putative surface proteins, of which 60% (2,663) were detected in our dataset (**Figure 3A**). We classified detected surface proteins into three specificity tiers across 42 cancer subtypes (n ≥10 per subtype): subtype-enriched (n=177), subtype-group-enriched (n=138), and broadly expressed (n=2,348) (**Figure S3A**). This classification assigned subtype- or group-level specificity to 39 of 42 cancer subtypes (**Figure S3B**) and recovered known selective targets. For example, GPNMB, targeted by glembatumumab vedotin and the CAR-T therapy GCAR1 (phase I in alveolar soft part sarcoma, NCT07297667), was among the most selectively elevated proteins in ASPS, cutaneous melanoma, uveal melanoma, and clear cell sarcoma. Consistent with the absence of a survival benefit when glembatumumab vedotin was tested in triple-negative breast cancer^40^, GPNMB abundance in our cohort was not elevated (**Figure S3C**). DCT, explored as a melanoma vaccine antigen^41^, showed similarly restricted abundance in cutaneous and mucosal melanoma subtypes (**Figure S3D**). TPBG, targeted across multiple indications (including BRCA, NSCLC and COADREAD) by the ADC ASN004 and the vaccine TroVax, was most highly expressed in GIST (**Figure S3E**). Beyond these confirmatory examples, the same analysis highlights biology-grounded discovery opportunities in rare cancers with limited treatment options: ANO1 in GIST, LDLR in ACC, ITLN1 in P(L)EMESO, CD36 in MRLS, A4GALT in RMS, RAPH1 and ACE in CHDM, METTL9 in UM, FGFRL1 in SACA, and SCARB1 in ACC and ASPS (**Figure S3F**, **Table S4**). Restricting the analysis to surface proteins with approved or active phase I–III drugs, 111 of 132 targets were quantified in the pan-cancer cohort with varying patient frequencies (**Figure 3B**, **Figure S4A**, **Table S5**). Applying a positivity threshold (z-score ≥1.5), 94% of patients were positive for at least one targetable surface protein (approved or in phase I–III; median=3, mean=3.9), and 38% were positive for at least one approved target (**Figure 3C**).

**Figure 3:**
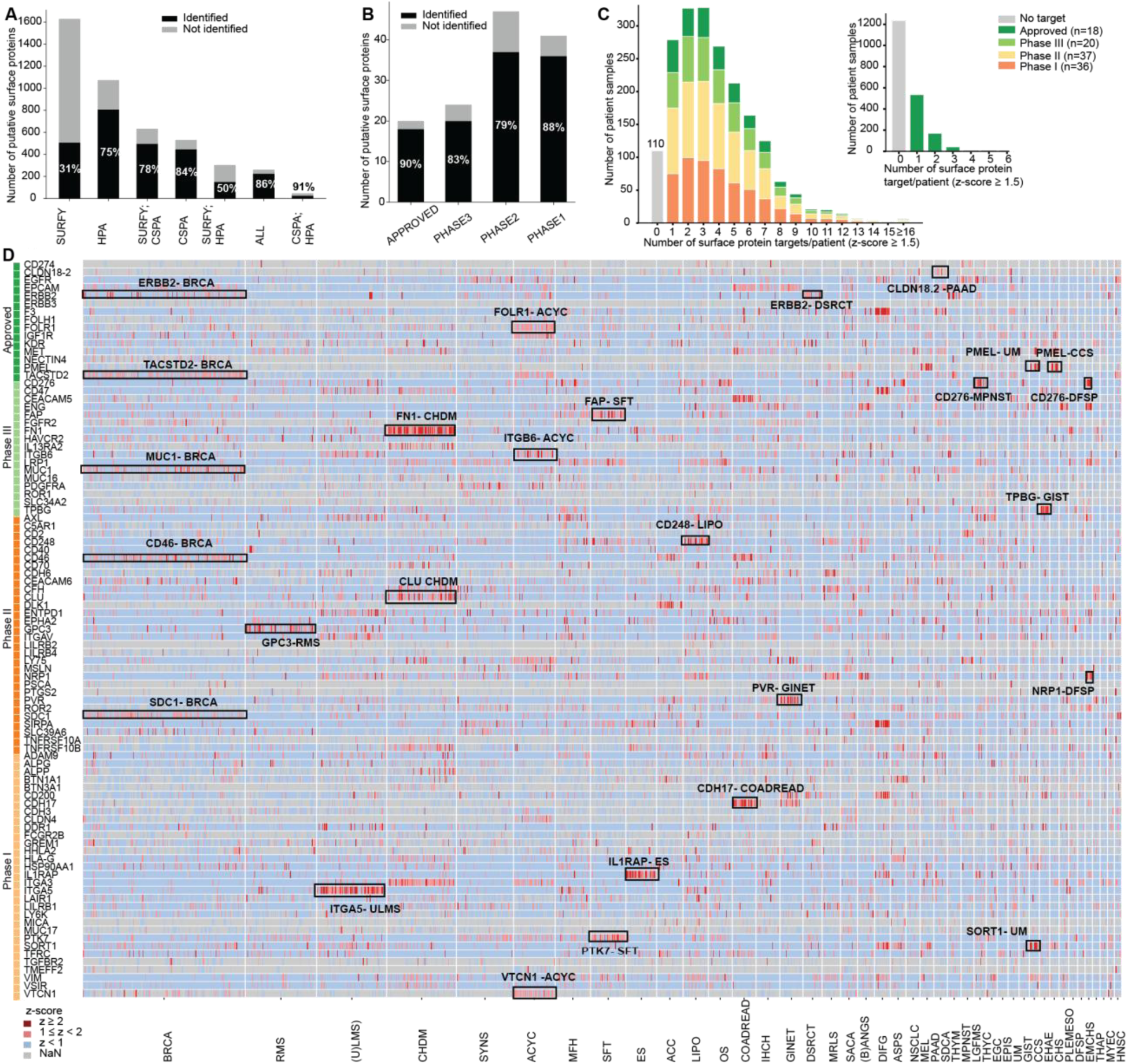
Expression landscape of druggable surface proteins across 42 cancer subtypes. (A) Number of identified (black) and unidentified (grey) proteins across three independent public surface-protein catalogs. (B) Numbers of identified (black) and unidentified (grey) surface proteins across clinical testing phases (Phase I–III, approved). (C) Number of surface protein targets per patient sample, colored by phase of clinical evaluation phase (green = approved, light green = Phase III, yellow = Phase II, orange = Phase I). Inset: Number of surface protein targets per patient sample for approved targets only. (D) Heatmap illustrating protein abundance (z-scored) for all surface targets in clinical evaluation or approved, across 42 cancer subtypes (n ≥ 10, NOS excluded). Rectangles in the heatmap depict selected surface target- cancer subtype pairs. *CSPA= Cell Surface Protein Atlas, HPA= Human Protein Atlas, SURFY= In silico human surfaceome*.

To illustrate the untapped potential for drug repurposing, we mapped expression of 93 quantified targets (n ≥200 patients) across 42 cancer subtypes. Proteins with approved or clinically evaluated drugs in a single indication frequently showed high expression across additional indications, including rare tumors with limited therapeutic options (**Figure 3D**). Trastuzumab deruxtecan (T-DXd), an antibody–drug conjugate targeting ERBB2, has demonstrated antitumor activity across a broad range of ERBB2-expressing solid tumors^42^ and was recently shown to be effective in desmoplastic small-round-cell-tumors (DSRCTs)^43^, an ultra-rare sarcoma with consistently high ERBB2 expression and limited treatment options (**Figure S4B**). FOLR1, the target of mirvetuximab soravtansine in advanced ovarian cancer, is highly expressed in adenoid cystic carcinoma (ACYC), a rare malignancy with no approved targeted therapy (**Figure S4C**). VTCN1 (B7-H4), explored in breast, ovarian, and endometrial cancer, is markedly elevated in ACYC as well (**Figure S4D**), identifying this histotype as a convergence point for multiple surface-directed strategies. The immune immune regulatory protein CD276 (B7-H3), under broad ADC investigation, showed particularly high expression in dermatofibrosarcoma protuberans (DFSP), malignant peripheral nerve sheath tumors (MPNST), and osteosarcoma (OS), three sarcoma subtypes sharing a pronounced unmet medical need (**Figure S4E**). Finally, TACSTD2 (TROP2), whose ADC sacituzumab govitecan delivers robust benefit in triple-negative breast cancer (TNBC), showed comparable abundance in cancers of unknown primary (CUPNOS) and ACYC (**Figure S4F**), extending its actionable landscape well beyond TNBC. While the systematic re-emergence of established targets across histotypes with limited treatment options provides a principled basis for repurposing approved agents beyond their original indications, the identification of additional proteins with restricted, high-abundance expression in rare and advanced cancers constitute tractable starting points for de novo drug development.

### A phosphoproteomic immune activity score captures functional states of ICI-responsive tumors

Immune checkpoint inhibition (ICI) has produced durable remissions in certain cancer types^44^, yet response rates across most remain modest (often ≤10%^45,46^). This dichotomy has made identifying predictive biomarkers a priority. Microsatellite instability-high (MSI-H), mismatch repair deficiency (dMMR) status^47^ and tumor mutational burden (TMB)^48,49^ currently act as surrogates for neoantigen load. However, CD8+ T-cell abundance estimated by transcriptomics outperformed TMB and also ranked highest among 36 neoantigen-, immune-, and checkpoint-related variables in predicting ICI response across 21 cancer types^50^. This suggests that the number and, potentially, activation state of tumor-infiltrating T-cells may be the more proximal determinants of therapeutic ICI benefit. T-cell activation is a phosphorylation-driven signaling event, downstream of the T-cell receptor (TCR) and checkpoint molecules, that genomics and transcriptomics cannot resolve^51,52^. We, therefore, propose a phosphoproteomic immune activity score (IAS) capturing T-cell activation-associated signaling to stratify patients for ICI therapy. IAS integrates three subscores (**Table S6**), each targeting a distinct layer of the cancer-immunity cycle. The antigen presentation score captures the MHC class I and II processing and presentation machinery, reflecting the tumor’s intrinsic capacity to generate and display neoantigens. The T-cell abundance score comprises proteins marking the presence and lineage identity of tumor-infiltrating lymphocytes, independent of activation state. The T-cell activity score reports on the functional, signaling-competent, state of infiltrating T-cells using (a total of 111) proteins, phosphoproteins, and p-peptides representing proximal TCR-signaling components and adaptors, transcriptional effectors, and markers of T-cell activation and exhaustion, **Figure S5A**). The IAS varied across cancer histologies, with overall higher scores in mesothelioma, melanoma, and carcinoma compared to sarcoma and neuroendocrine neoplasms (**Figure S5B).** This mirrors established clinical response patterns: melanoma and mesothelioma are among the histologies with the highest rates of durable ICI benefit^53^, whereas neuroendocrine neoplasms show consistently low response rates across ICI regimens^54^. Sarcomas as a group have long been considered “immune-cold,” yet are immunologically heterogeneous across subtypes^55^, a pattern reflected by intermediate and broadly distributed scores observed here. However, within the sarcoma group, alveolar soft part sarcoma (ASPS), angiosarcoma (ANGS), and epithelioid sarcoma (EPIS) showed the greatest proportion of IAS-high patients and the highest median scores (**Figure 4A, Table S7**). This aligns with known ICI responsiveness in ASPS^56^ and ANGS^57^, and, to a lesser extent, EPIS^58^, where activity is largely reported in basket trials and case reports. Correlating the IAS with established genomic and transcriptomic ICI biomarkers, we observed only moderate associations with TMB and PD-L1 mRNA (**Figures 4C and S5C**), suggesting the score captures complementary information. PLEMESO and ASPS exemplify this disconnect, responding well to ICI despite low TMB and inconsistent PD-L1 expression^53,56,59,60^. Consistently, both subtypes showed elevated IAS relative to the pan-cancer median in our cohort while TMB and PD-L1 mRNA were not elevated (**Figure S5D**), illustrating that the IAS reflects functional immune activation independently of neoantigen load and checkpoint-ligand abundance. This pattern was further apparent when comparing IAS z-scores across subtypes with high versus low ICI response rates (**Figure 4B**). In an exploratory pan-cancer cohort treated with ICIs (n = 31, 20 cancer subtypes), IAS correlated positively with progression-free survival (PFS, HR = 0.71, 95% CI 0.53–0.97, p = 0.03, **Figures S5E and S5F, Table S8**). Stratification by a pre-specified median IAS split reinforced this finding, with IAS-high patients achieving a median PFS of 17.0 months compared to 3.3 months in IAS-low patients (HR = 0.38, 95% CI 0.14–0.98; log-rank p = 0.040; **Figure 4D**) and a significantly higher disease control rate (89% vs. 40%; Fisher’s exact p = 0.020, Wilcoxon rank-sum p = 0.016; **Figure 4E**). Together, these results suggest that IAS may serve as a pan-cancer predictive biomarker for ICI response, warranting prospective validation in larger, histology-stratified cohorts.

**Figure 4:**
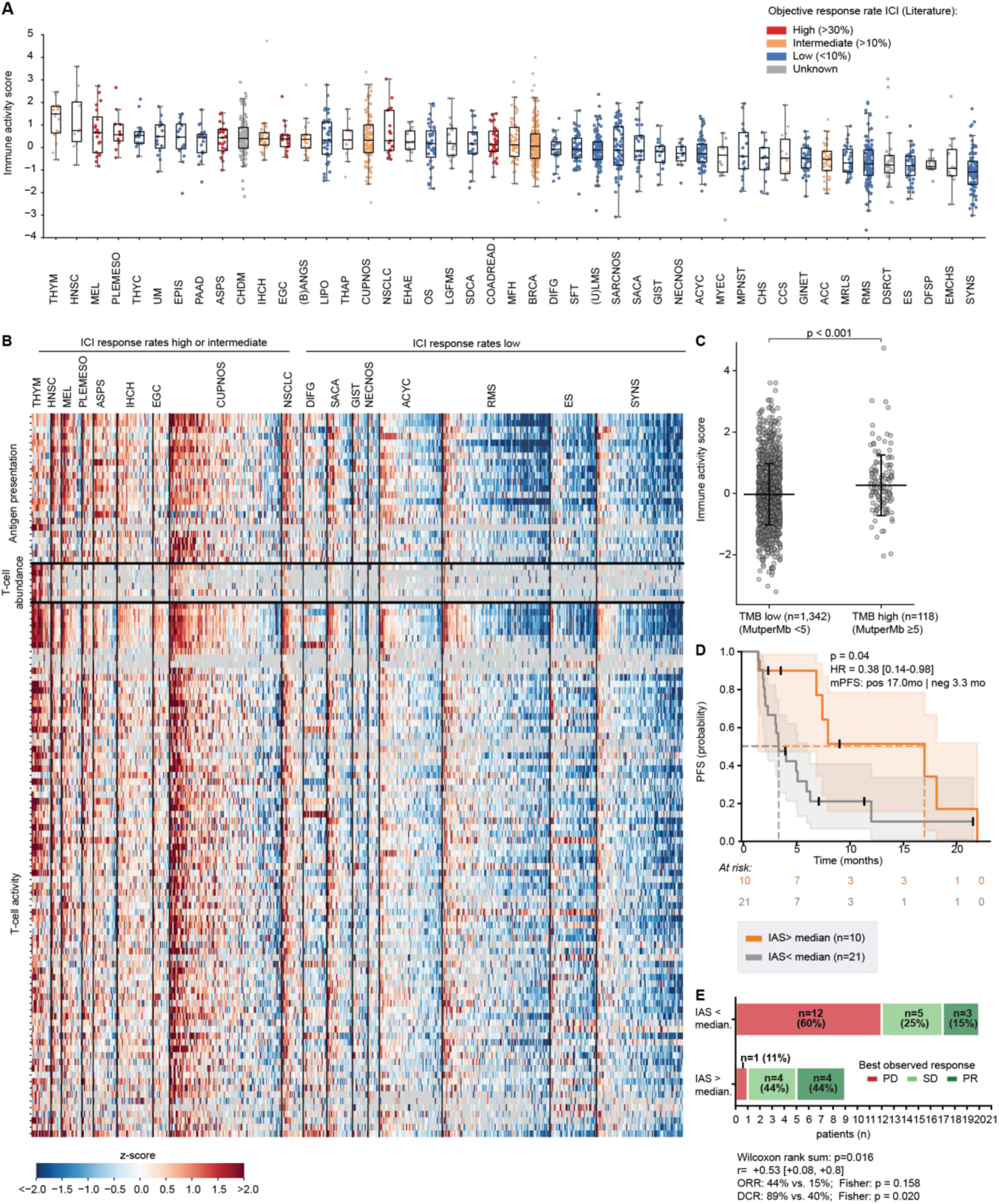
Phosphoproteomics-derived immune activity score. (A) Box plot illustrating the immune activity score across 44 cancer subtypes (n ≥ 10, sorted by highest median, left to right), colored by clinical trial-reported objective response rate (ORR) to immune checkpoint inhibition (ICI): red = high ORR (>30%), orange = intermediate ORR (10–30%), blue = low ORR (<10%), grey = unknown ORR (no clinical trial data). (B) Heatmap depicting z-scored protein abundance, p-peptide abundance, and protein phosphorylation scores across the three immune activity subscores, for patient samples from selected cancer subtypes with high, intermediate, or low ORR to ICI. (C) Comparison of the immune activity score in TMB low (MutperMb <5) and TMB high (MutperMB >=5) tumor samples. The horizontal bar indicates the median; error bars denote ±1 standard deviation. (D) Kaplan–Meier analysis of progression-free survival under ICI therapy stratified by immune activation score. PFS estimated by the Kaplan–Meier method in IAS-positive (orange; immune activation score > −0.0035; n = 10) and IAS-negative (grey; score ≤ −0.0035; n = 21) patients. Tick marks indicate censored observations; shaded areas represent 95% pointwise confidence intervals. Numbers at risk are shown below the plot at 5-month intervals. (E) Horizontal stacked bar chart showing the distribution of best observed response (BOR) within each IAS group (IAS-positive n = 9; IAS-negative n = 20; NA excluded).

### Pan-cancer landscape of clinically relevant kinase activities

We developed Tumor Proteome Activity Status (TOPAS) scores to infer the activity of 46 kinases relevant to cancer biology (Methods, see accompanying manuscript) and applied these to 32 cancer subtypes (n≥10 patient samples, **Figure 5, Table S9**). The TOPAS panel spans RTKs, mitogenic MAPKs, stress MAPKs, PI3K-AKT-mTOR pathway members, cell cycle and transcriptional kinases as well as DNA damage kinases, many of which represent established or emerging therapeutic targets. The resulting pan-cancer kinase activity landscape recapitulated canonical genomic RTK driver associations. EGFR activity was elevated in subsets of non-small cell lung carcinoma (NSCLC) and diffuse glioma (DIFG), consistent with the high frequency of activating EGFR alterations in these entities. KIT activity was prominent in gastrointestinal stromal tumors (GIST) and thyroid carcinoma (THYC), and PDGFRB in DFSP, reflecting that KIT mutations and COL1A1–PDGFB fusions define these respective tumor types,.

**Figure 5:**
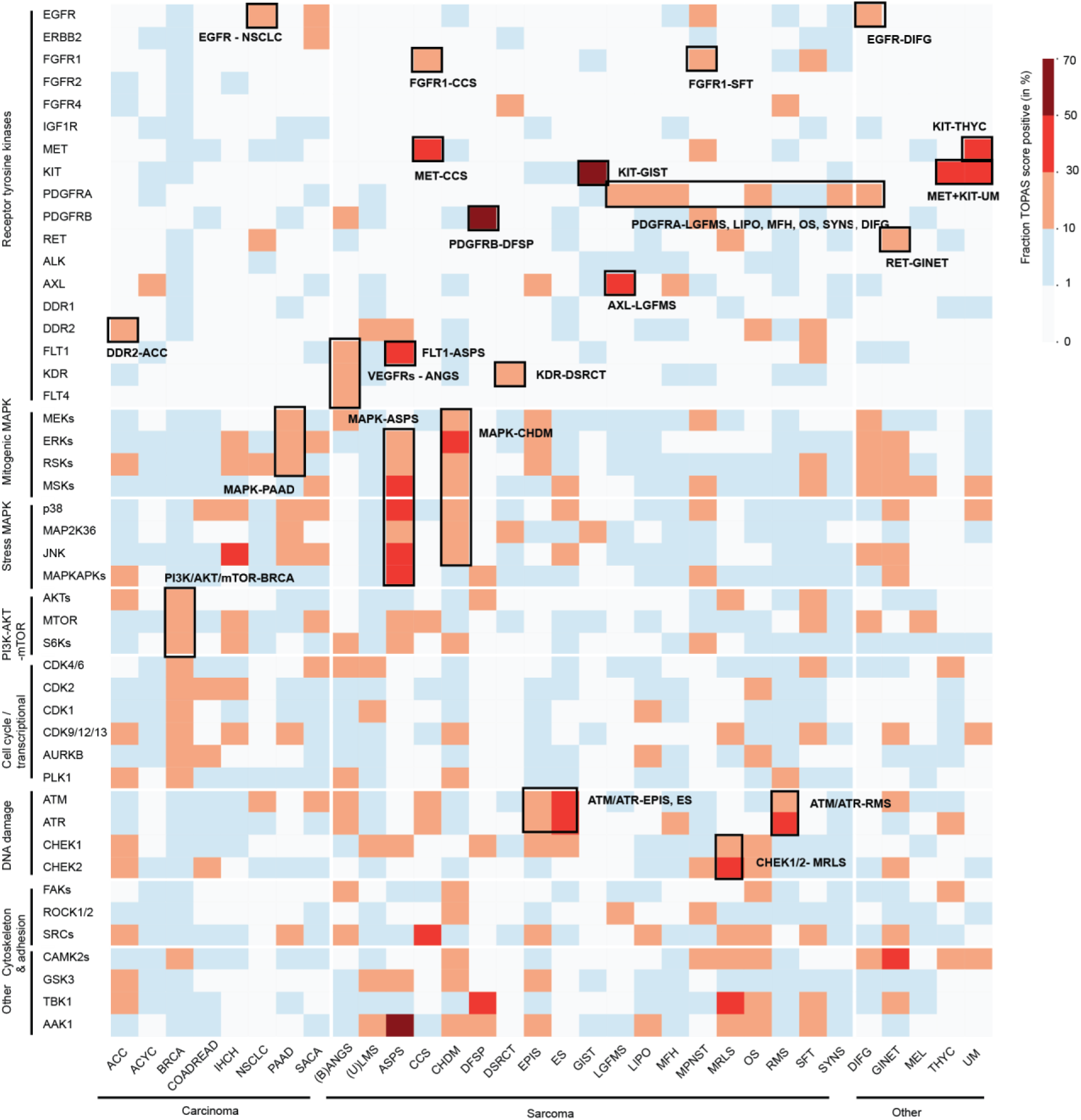
Pan-cancer landscape of clinically relevant kinase activities. Heatmap depicting TOPAS-positivity (fraction of samples with TOPAS score ≥ 2) for 46 kinases spanning different kinase categories/pathways, across 32 cancer subtypes (n ≥ 10; samples with limited sample quality excluded), sorted by broad histology (carcinoma, sarcoma, other). Rectangles highlight selected kinase–cancer subtype pairs.

Beyond these established associations, the landscape revealed a broad set of RTK activities not readily explained by known genomic alterations, pointing to non-genomic mechanisms of kinase activation and potentially novel therapeutic vulnerabilities. VEGFR family members FLT1, KDR, and FLT4 were collectively elevated in breast and non-breast angiosarcomas (**Figure S6A**), reflecting the vascular lineage of these tumors rather than recurrent genomic alteration of the receptors, which occurs in only a minority of cases. High FLT1 activity was also detected in alveolar soft part sarcoma (ASPS), and high KDR in a subgroup of desmoplastic small round cell tumors (DSRCT). MET and KIT co-activation emerged in uveal melanoma (UM), while MET together with FGFR1 was prominent in clear cell sarcoma (CCS), suggesting RTK co-activation as a shared feature of these rare melanocytic tumors (**Figure S6B and S6C**). PDGFRA activity was broadly elevated across multiple sarcoma subtypes and DIFG (**Figure S6D**). Additional subtype-restricted activities included FGFR1 in solitary fibrous tumor (SFT), RET in gastrointestinal neuroendocrine tumors (GINET), DDR2 in adrenocortical carcinoma (ACC), and AXL in low-grade fibromyxoid sarcoma (LGFMS).

### RTK activity and genomic status are frequently discordant in advanced cancer

Somatic RTK alterations are classified as oncogenic or likely oncogenic in resources such as OncoKB and CIViC based on hotspot mutation recurrence, functional evidence, and clinical data^61,62^. These classifications are derived from well-characterized, high-incidence tumor types, but their ability to predict RTK activity across diverse histological contexts is unclear. This is apparent in our cohort, which spans rare malignancies with sprase histotype-specific evidence^63^ and is populated by advanced, heavily pre-treated patients whose tumors carry complex evolutionary histories shaped by therapeutic selection. Under these conditions, clonal dynamics can decouple detectable genomic alterations from active RTK signaling, because subclonal or therapy-enriched events may no longer function as dominant oncogenic drivers^64^. We, therefore, examined how RTK TOPAS scores relate to matched oncogenic and likely oncogenic RTK alterations to assess the actionability implied by genomic findings. Of 306 patients with genomic alterations classified as oncogenic (n=182) or likely oncogenic (n=124) across 18 RTKs, only 72 (24%) were associated with an elevated RTK TOPAS ≥2 (**Figure 6A**). Recall was twice as high for oncogenic as for likely oncogenic events (54/182, 30% vs. 18/124, 15%), consistent with the expected higher functional impact of confirmed driver alterations. Recall rates also varied substantially across RTKs. For oncogenic alterations, ALK (3/4, 75%), ERBB2 (21/38, 55%), PDGFRA (9/18, 50%), and EGFR (11/29, 40%) showed robust recall, whereas FGFR1 (2/43, 5%), FGFR4 (0/8, 0%), and KDR (1/14, 7%) showed poor recall, indicating that genomic driver status alone is insufficient to predict functional kinase activation. For gene amplifications, recall ranged from 68% for ERBB2 (21/31) to below 10% for KDR, KIT, DDR2, FGFR1, FGFR4, and ALK (**Figure 6B**). Notably, ERBB2, the only current RTK with a tumor-agnostic level 1 biomarker designation for strong protein overexpression (IHC 3+), showed good concordance between ERBB2 copy number and TOPAS scores across cancer subtypes, with recall reaching 91% at total copy numbers (TCN) of ≥20 (**Figure S6E**). This reflects the potent functional consequence of ERBB2 amplification and suggests that raising the threshold for amplification calling may improve the specificity of actionability predictions.

**Figure 6:**
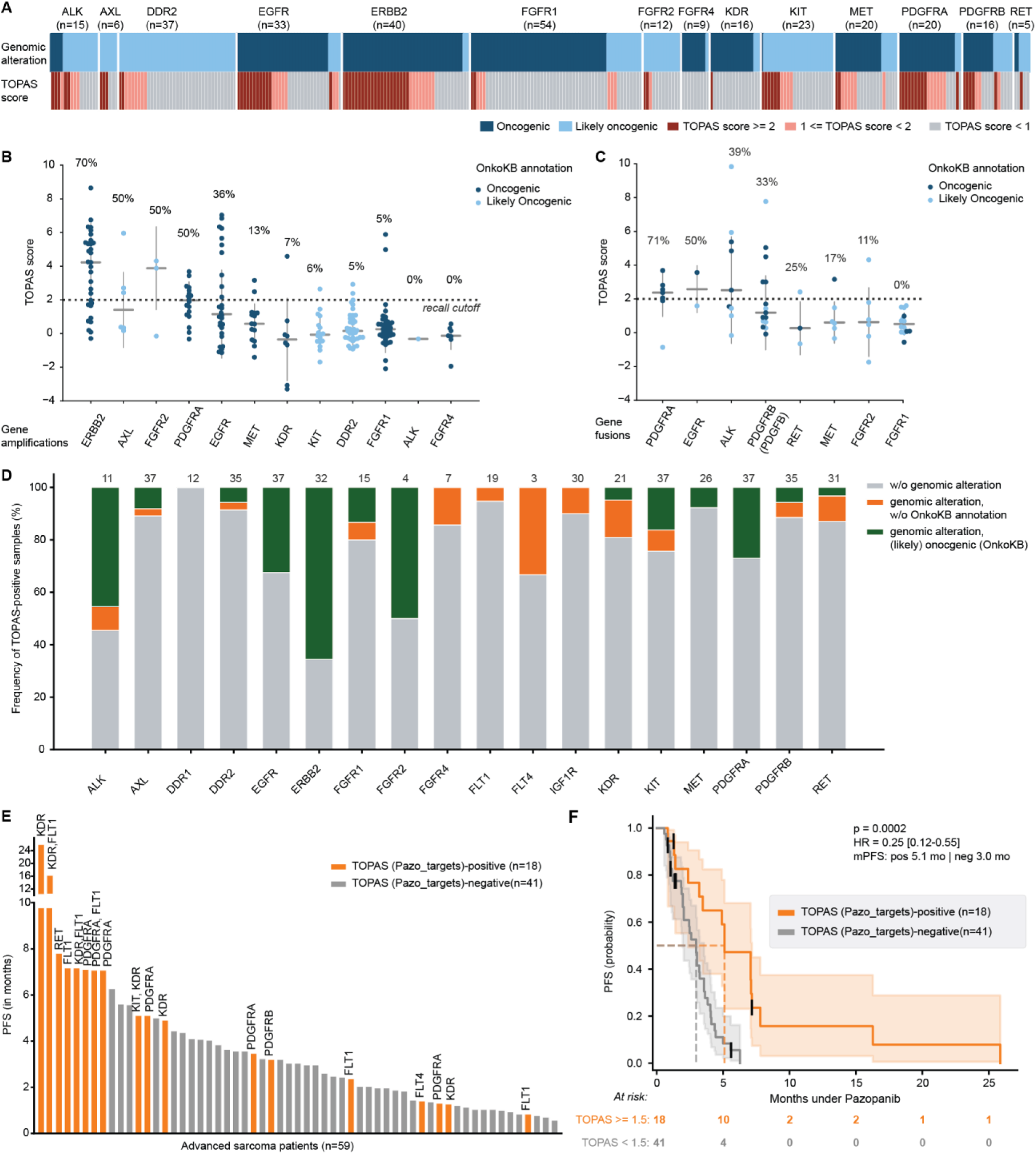
RTK activity and genomic status in advanced cancers. (A) Recall of OnkoKB-annotated oncogenic (blue) or likely oncogenic (light blue) genomic RTK alteration by TOPAS positivity (≥2 = red, ≥1 light red, <1= grey) for 14 (out of 18, filtered for ≥5 annotated genomic alterations) RTKs. (B) Recall of RTK gene amplification events by positive TOPAS score (≥2). Fraction of recall is indicated in the plot for 12 RTKs (n≥5 samples with gene amplification). (C) Recall RTK gene fusion events by positive TOPAS score (≥2). Fraction of recall is indicated in the plot for 8 RTKs (n≥ 5 samples with gene fusion). For PDGFRB, PDGFB fusions are included in the analysis. (D) Stacked bar plot per RTK, indicating the fraction of TOPAS-positive samples with OnkoKB-annotated genomic RTK alteration (green), with genomic alteration (not annotated, orange) and without any genomic alteration affecting that specific RTK. Total number of TOPAS-positive samples is indicated in the plot above the respective bar. (E) Ranked PFS (in months) of 59 advanced sarcoma patients colored by TOPAS-positivity (≥1.5, orange) for one out of 7 pazopanib targets (FLT1, KDR, FLT4, PDGFRA, PDGFRB, KIT, RET). (F) Kaplan-Meier curves depicting progression-free survival (PFS) probability under pazopanib therapy in a cohort of advanced sarcoma patients (n = 59) stratified by the maximum TOPAS score across pazopanib target kinases. Patients were split into TOPAS-positive and TOPAS-negative groups using a score threshold of ≥ 1.5 vs. < 1.5 (n = 18/41). PFS was defined as the time from therapy initiation to progressive disease (PD) or death. Patients discontinuing therapy due to toxicity (n = 4) or for unknown reasons (n = 3) were censored and are indicated by vertical bars (|). Shaded areas = 95% confidence intervals (Greenwood’s formula). Dashed lines indicate median PFS. Numbers at risk shown below at 5-month intervals. HR with 95% CI from univariable Cox proportional hazards model; TOPAS-negative group as reference. Orange: TOPAS-positive; grey: TOPAS-negative.

The overall low recall for most RTKs underscores the context-dependence of RTK alteration functional impact and highlights the value of orthogonal functional measurement for interpreting the actionability of genomic findings. The same pattern held for genomic fusion events. Across 73 patient samples with RTK gene fusions (10 RTKs) and annotated as oncogenic (n=28) or likely oncogenic (n=45), 19 (26%) were recalled by a TOPAS score ≥2 (**Figure 6C**). Again, oncogenic fusions were recalled at higher rates than likely oncogenic fusions (14/28, 50% vs. 5/45, 11%). Recall was highest for ALK (3/4, 75%) and PDGFRA fusions (5/6, 83%), consistent with the well-established functional consequence of fusion-driven kinase activation for these receptors. In contrast, FGFR1 fusions, the most frequent RTK fusion event in this cohort (n=15), showed no recall (0/15, 0%), pointing to a consistent disconnect between FGFR1 genomic alteration and phosphoproteomic evidence of kinase activation. This aligns with clinical observations in which FGFR1 amplification has repeatedly failed as a predictive biomarker for FGFR inhibitor response^65^, and with mechanistic evidence that FGFR1 amplicons are structurally heterogeneous and may not confer functional kinase dependence^66,67^. A further illustration of this complexity came from the COL1A1-PDGFB fusion in dermatofibrosarcoma protuberans (DFSP), a pathognomonic rearrangement of DFSP: approximately half of affected cases failed to produce outlier PDGFRB kinase activity, coinciding with absent or reduced expression of the receptor or its ligand (**Figure S6F**). Even a well-characterized oncogenic fusion within a defined histologic subtype is, therefore, insufficient to guarantee functional RTK activation.

### RTK activity in the absence of genomic alterations informs patient stratification and therapy selection

RTK TOPAS scoring identified a substantial population of tumors with increased RTK signaling that would have been entirely missed by genomic profiling: 34% (354/1,048) of samples in the cohort showed at least one outlier RTK TOPAS score. Of those, 70% (248/354) lacked an oncogenic or likely oncogenic RTK annotation, and 197 of these lacked any genomic RTK alteration (**Figure 6D**). This indicates that non-genomic mechanisms of RTK activation, including transcriptional upregulation, post-translational regulation, paracrine or autocrine ligand stimulation, and epigenetic reprogramming, are widespread yet systematically under-detected by sequencing-based approaches. These cases represent a reservoir of potentially targetable RTK dependencies that current “genomics-first” paradigms would fail to prioritize. Unlike many carcinomas where oncogenic RTK alterations define molecular subtypes, the clinical relevance of non-genomic RTK activity is particularly apparent in sarcoma, which are predominantly driven by chromosomal translocations and complex karyotype instability^68,69^. While sarcomas are frequently RTK-TOPAS positive (37%), most (79%) of these had no accompanying (likely) oncogenic genomic RTK alteration (**Figure S6G**). Despite the absence of a genomic mechanism, the multi-RTK inhibitor pazopanib improved PFS in advanced soft-tissue sarcoma, leading to regulatory approval^70^. This suggests that kinase-activity outlier status may capture biologically and clinically meaningful RTK dependence irrespective of genomic annotation. To investigate the utility of TOPAS scoring for therapy response stratification, we analyzed 59 advanced sarcoma patients for whom pre-treatment biopsies were available. Patients showed a wide range of PFS intervals (17-776 days; **Table S10**) and we observed a clear relationship between longer PFS and elevated TOPAS scores for pazopanib targets (FLT1, KDR, FLT4, PDGFRA/B, RET, KIT) (**Figure 6E, Figure S6H**). Kaplan-Meier analysis showed a significant PFS benefit in the 18 patients with elevated TOPAS scores (≥1.5) for at least one pazopanib target (mPFS 5.1 months) compared to the 41 patients with lower scores (mPFS 2.5 months; HR = 0.25, 95% CI 0.12–0.52; p = 8×10⁻⁵; **Figure 6F**). This stratification was robust to TOPAS score threshold choice: both the conservative TOPAS score ≥2 cutoff and an unbiased median split (≥ 1.02) resulted in significant PFS differences (**Figures S6I and S6J**). Together, these results establish that pan-cancer kinase-activity profiling reveals a layer of RTK biology that is orthogonal to, and substantially broader than, what can be inferred from genomic data, with direct implications for patient stratification and treatment selection.

### Interpreting intracellular kinase activity requires lineage, genomic, and pathway context

Unlike RTKs, whose oncogenic activation is often a potent, lineage-spanning event, intracellular kinases present two distinct challenges for activity inference. First, they often function as convergent signaling nodes that integrate inputs from multiple upstream pathways. Even a strong oncogenic driver, whether a kinase-activating mutation within the pathway itself or an upstream RTK alteration, may only partially contribute to total kinase activity, with the remainder reflecting the broader signaling state of the cell. In advanced and heavily pre-treated tumors, where pathway remodeling and adaptive rewiring are common, such signal dilution is likely further compounded. Second, the baseline activity of intracellular kinases varies substantially across cancer subtypes. For example, ERK activity (**Figure 7A)** and AKT activity (**Figure 7B**) reflect cell-type-specific differences in pathway wiring, co-factor availability, and transcriptional programs that are independent of oncogenic activation. Both challenges for kinase activity scoring become apparent when attempting to associate oncogenic alterations in the RAS-RAF-MEK-ERK axis with ERK and RSK TOPAS scores (**Figures 7C and S7A**), or, analogously, in the PI3K-AKT axis with AKT and S6K scores (**Figures 7D and S7B**). In both cases, tumors harboring oncogenic alterations show statistically higher activity scores than those without. However, effect sizes are small, and distributions broadly overlap, precluding meaningful patient-level interpretation. Breaking this analysis down to subtypes provided more clarity. For instance, RAS–RAF alterations elevate ERK TOPAS scores in pancreatic (PAAD) and colorectal (COADREAD) cancer (**Figure S7C**), where RAS-ERK signaling is the established dominant oncogenic axis^71,72^. In breast cancer (BRCA), however, RAS-RAF alterations do not increase ERK TOPAS scores. Similarly, breast carcinomas, salivary carcinomas (SACA), and myxoid liposarcoma (MRLS) harboring oncogenic alterations in any PI3K or AKT isoform trended toward high AKT activity, while this association was absent in several other subtypes, including adenoid cystic carcinoma (ACYC) and leiomyosarcoma (LMS, uterine and non-uterine; **Figure S7D**). A complementary pattern emerged when stratifying by alteration class. SNVs and gene fusions affecting RAF, RAS, and MEK isoforms were associated with more robust and consistent ERK activity elevation than amplifications (**Figure S7D).** Similarly, SNVs in PI3K or AKT isoforms were associated with more robust AKT activity elevation than fusions, amplifications or deletions of regulatory PI3K isoforms. This distinction reflects a fundamental difference in activation mechanism: point mutations constitutively lock the kinase in its active state, whereas amplifications increase protein abundance without bypassing normal regulatory checkpoints, making pathway output dependent on the broader signaling state of the cell. The lineage- and alteration-context dependence is well illustrated in GIST: KIT-activating mutations account for ∼75–80% of cases and are rarely accompanied by recurrent RAS, RAF, or other MAPK- or AKT-activating co-mutations^73,74^, providing an unusually clean oncogenic background for attributing ERK and AKT activity to KIT. Consistently, KIT-mutant tumors showed clearly elevated mitogenic MAPK and AKT signaling relative to KIT-wild-type counterparts, while tumors sampled under therapy showed activity levels comparable to the KIT-wild-type group, confirming on-target drug activity (**Figure 7E**). Yet, because GIST operates at a characteristically low baseline for both pathways, the absolute z-scores of even KIT-mutant tumors were overall low within the pan-cancer range, in fact falling below any threshold for calling aberrant activity and obscuring what is, within the lineage, a robust oncogenic signal. Together, these observations indicate that intracellular kinase activity is not a direct readout of oncogenic alteration status, but an integrated signal shaped by lineage, alteration type, and broader pathway context, a complexity that lineage-aware normalization can help account for but not fully resolve.

**Figure 7:**
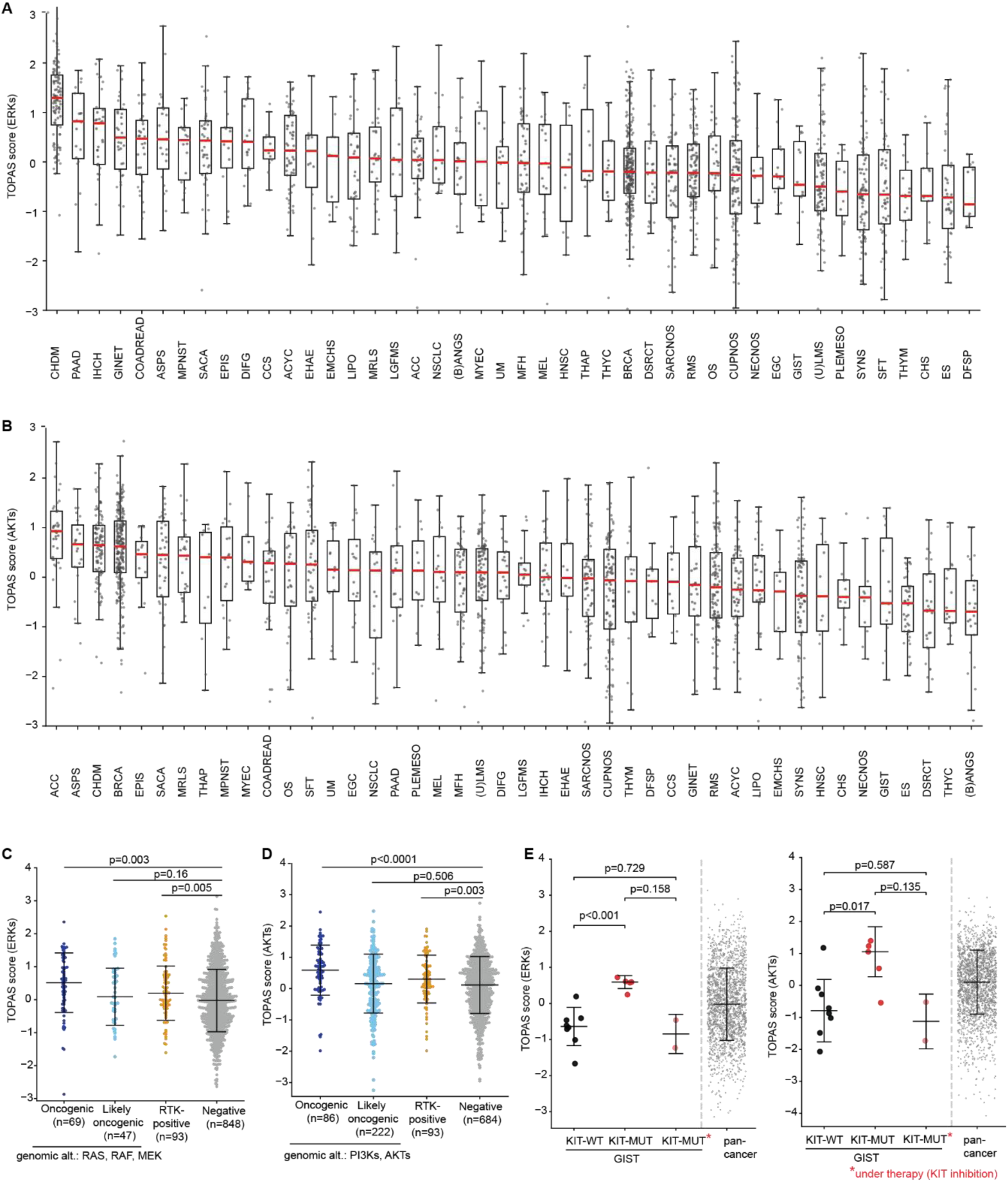
Subtype-resolved landscape of intracellular growth kinase activities. (A, B) ERKs (A) or AKTs (B) TOPAS scores across 44 cancer subtypes (n ≥ 10), ranked by median TOPAS score (highest to lowest). (C) ERK TOPAS score shown for patient samples grouped by genomic alteration status in RAS/RAF/MEK or RTK-positivity: at least one oncogenic (blue) or likely oncogenic (light blue) alteration in one of the RAS/RAF/MEK isoforms, or RTK-positive (RTK TOPAS score ≥ 2 and a genomic RTK alteration; orange), vs. negative (none of the three criteria; grey). (D) AKTs TOPAS score shown for patient samples grouped by genomic alteration status in PI3K/AKT or RTK-positivity: at least one oncogenic (blue) or likely oncogenic (light blue) alteration in one of the PI3K or AKT isoforms, or RTK-positive (RTK TOPAS score ≥ 2 and a genomic RTK alteration; red), vs. negative (none of the three criteria; grey). (E) ERKs (left) or AKTs (right) TOPAS score shown for gastrointestinal stromal tumor (GIST) patients grouped by oncogenic KIT mutation (red, orange; orange = sampled under KIT-inhibitor therapy) or KIT-WT (black), compared to the pan-cancer cohort (grey). For all swarm plots: Each dot represents one tumor sample. The horizontal bar indicates the median; error bars denote ±1 standard deviation. Statistical comparisons between each group and the negative reference group were performed using a two-sided Welch’s t-test; p-values are shown above brackets.

### A lean phosphoproteomic biomarker stratifies chordoma patients for EGFR-targeted therapy

Chordoma (CHDM) is a rare notochordal tumor most commonly occurring at the skull base, mobile spine, and sacrum. Local recurrence rates are high (>50%), and many advanced patients develop metastatic disease^75^. For these patients, treatment options are limited, and no FDA-approved therapy currently exist. Several studies have reported aberrant activity of PDGFRB, KIT, MET, and EGFR^76,77^. However, molecularly unstratified clinical trials with KIs have shown only modest benefit^78–80^. Anecdotal responses to EGFR inhibitors have been reported^81,82^ and a phase II trial of the EGFR kinase inhibitor afatinib (NCT03083678) reported a 12-month PFS rate of 40%^83^. To evaluate whether the TOPAS approach may help select CHDM patients for therapy, we analyzed the RTK landscape in 102 tumor specimens from 86 chordoma patients. Aberrant RTK activity (TOPAS score ≥2) was detected in 19% (19/102) of samples covering seven RTKs, with KDR, PDGFRB and EGFR being the most prevalent (**Figure 8A**). Combined genome, transcriptome, and phosphoproteome analyses revealed that abnormal EGFR activity was detectable exclusively by the phosphoproteome (**Figure S8A**). To investigate whether high EGFR TOPAS scores may be indicative of therapy response to EGFR inhibitors, we measured phosphoproteomes and phenotypic responses of 14 CHDM cell lines and 11 CHDM patient-derived xenograft (PDX) models to afatinib (**Figures 8B,8C andS8B**). Samples were classified as drug responders (EC50 ≤50 nM for cell lines; treatment arm/control arm tumor volume (T/C TV) ratio <0.4 for PDX models) or non-responders (EC50 ≥1,000 nM for cell lines; T/C TV ratio ≥0.4 for PDX models). Although there were more cases with elevated EGFR TOPAS scores (EGFR+) in the responder group, the TOPAS score alone was insufficient to predict responses (**Fig. 8D**). This indicates that EGFR overactivity is neither necessary nor sufficient for EGFR dependency. Differential protein expression analysis of EGFR+ responder vs non-responder models revealed high levels of interferon (IFN)-induced proteins (ISGs, e.g. MX1) in the non-responder group (**Figure S8C**). These proteins have also been found to be inversely correlated with response to EGFR inhibitors in lung cancer^84^ and are associated with resistance to EGFR inhibitors in chordoma^85^(**Figure 8D**). Similarly, differential protein expression analysis of EGFR TOPAS-low (EGFR-) responders and non-responders showed that the abundance of TUSC3, a protein glycosyltransferase shown to reduce EGFR phosphorylation^86^, was significantly elevated in responder models. Elevated TUSC3 potentially suppresses EGFR phosphorylation sufficiently to place these tumors below the EGFR TOPAS score reporting threshold, classifying these patients as EGFR- despite underlying EGFR dependency (**Figures 8D and S8D**). By combining EGFR TOPAS scores with either MX1 abundance (for EGFR+ models) or TUSC3 abundance (for EGFR- models), 24/25 chordoma models were correctly classified (**Figure 8E**). To assess whether this preclinical stratification principle is consistent with clinical outcomes, we applied the combined biomarker retrospectively to the seven patients in our chordoma cohort who had received EGFR inhibitors (some in combination with bevacizumab), classifying them as clinical responders (>12 months therapy duration, n=5) or non-responders (≤6 months therapy duration, n=2) (**Figure 8F**). By assigning each chordoma sample as a responder (n=54) or non-responder (n=48) on the basis of the combined EGFR TOPAS score and MX1/TUSC3 biomarker, the predicted class was concordant with clinical outcome in 6/7 treated patients (**Figure 8G**). While the retrospective cohort is too small for definitive conclusions, concordance between preclinical and clinical classifications establishes a proof-of-stratification principle, supporting prospective evaluation in ongoing EGFR inhibitor trials in chordoma.

**Figure 8:**
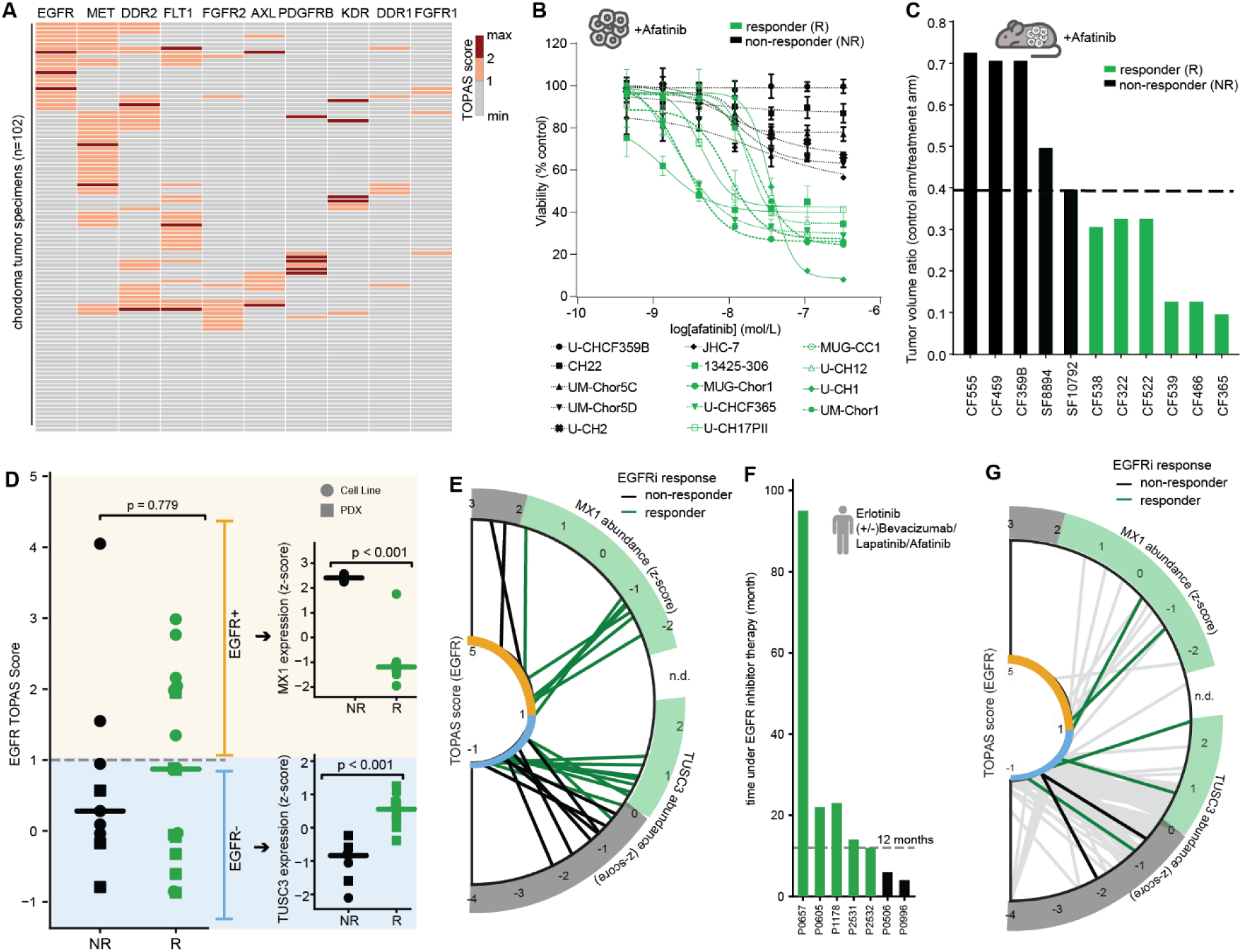
Stratification of chordoma patients for EGFR-targeted therapy. (A) Activity landscape in advanced chordoma patients (n=102 samples; 86 patients) for 10 RTKs. (only scores with n≥ 5 samples with TOPAS score ≥1 displayed). Samples are clustered by the respective RTK TOPAS score (≥1). (B) CellTiter-Glo proliferation assays of chordoma cell lines treated with afatinib (biological replicates = 2-3/cell line). Sensitive cell lines (EC50 <50nM) are colored in green and resistant cell lines (EC50>1000nM) in black. (C) Afatinib response in patient-derived xenograft (PDX) models (5-7 mice/arm). Graphs show the ratio of mean tumor volume in the therapy arm (20mg/kg afatinib) over the control arm at day of assessment (control arm >1500mm³ or at day 42). Models with a T/C ratio ≤ 0.5 are classified as responders and are colored in green. (D) Swarm plot illustrating EGFR TOPAS scores in the non-responder (n=9, black) and responder group (n=16, green) across chordoma cell lines (dot) and PDX models (rectangle). Differential abundance of MX1 in the EGFR TOPAS positive (z-score ≥1) responder and non-responder phenotypes. Differential abundance of TUSC3 in the EGFR TOPAS negative (z-score <1) responder and non-responder phenotypes. (E) Parallel coordinate plot using EGFR TOPAS socres and either MX1 protein expression (z-score) in case of EGFR TOPAS positivity or TUSC3 protein expression (z-score) in case of EGFR TOPAS negativity for chordoma model (cell line and PDX) stratification. Lines connecting coordinates are colored in green (responder phenotype) or black (non-responder phenotype). (F) Bar plot indicating the time under EGFR inhibition (erlotinib+bevacizumab, lapatinib or afatinib) in 7 patients. Patients with a therapy duration > 12 months (dashed grey line) are classified as responders and are colored in green. (G) Parallel coordinate plot using EGFR TOPAS scores and either MX1 protein expression (z-score) in case of EGFR TOPAS positivity or TUSC3 protein expression (z-score) in case of EGFR TOPAS negativity for chordoma patient stratification. Lines connecting coordinates for patients with clinical response are colored in green, for patients without response in black, and for patients without EGFR-therapy follow-up in grey.

## Discussion

While genome-focused MTBs are now common in major academic cancer centers and precision-oncology programs, we are the first to incorporate comprehensive phosphoproteome profiling into a prospective, multi-center, real-world pan-cancer setting. Because most oncogenic mechanisms lead to malfunctioning proteins that are the targets of most cancer drugs, this constitutes an important conceptual advance. Processing >2,500 cases over five years demonstrates sustained technical feasibility, and the identification of >8,000 proteins and >25,000 p-peptides per patient ensured broad coverage of actionable cancer-associated genes and precision oncology drug targets. The data proved valuable along multiple lines: the proteome expression data enabled training the PROdict classifier that predicts cancer subtypes even in a metastatic context. The cancer surface proteome highlights many opportunities for repurposing ADCs, CAR-T therapies, and other cell-surface-targeted modalities. The phosphoproteome enabled scoring of kinase and immune activity in tumor tissue, addressing important diagnostic gaps that DNA- and RNA-level data cannot fill. Proteogenomic analysis of >1,000 pan-cancer cases repeatedly showed added value of the cancer phosphoproteome beyond the (altered or unaltered) genome. For instance, in advanced and heavily pretreated tumors, diverse genomic lesions, their combinations, or non-genomic mechanisms converge on a limited number of central signaling nodes that phosphoproteomics can directly measure, providing functional readouts that cannot be inferred from genome sequencing. In rare cancers, this is particularly consequential because oncogenicity annotations are almost exclusively derived from common cancer types and transferred to rare entities that cannot be prospectively tested at scale. Phosphoproteomics can often resolve this ambiguity by reading out whether a transferred biomarker drives any signaling in the histological context of the individual patient. Together, the new data layers inform MTB recommendations for nearly every patient.

The current work complements and extends other cancer proteome profiling resources such as CPTAC and the pan-cancer proteome atlas^19,31,84^. CPTAC (∼6,000 cases from 39 cancer types) established that phosphoproteomics can reveal kinase-driven biology in the absence of established genomic drivers. The pan-cancer atlas (∼1,000 cases from 22 cancer types) showed that cross-continent expression proteomics data integration is feasible. Both resources were built retrospectively from treatment-naive resections of common cancers, with workflows and data pipelines optimized for cohort-level analysis. Our pan-cancer cohort (∼2,000 cases from 194 cancer types) is prospective and dominated by rare and heavily pretreated tumors with complex evolutionary histories for whom the medical need is greatest. In addition, our approach delivers quality-controlled results in time for MTB discussions of individual patients and systematically benefits from the increasing discriminating power of an ever-growing population of pan-cancer reference phosphoproteomes. This enables both patient-centric interpretation and cohort-level discovery for use today and in the future.

Despite this progress, the study has limitations. For instance, PROdict currently covers only 26 of 194 cancer subtypes, TOPAS scores have been developed for just 46 kinases and the broader clinical impact of phosphoproteome-guided MTB recommendations cannot yet be systematically and objectively evaluated because the presented clinical analyses on explaining pazopanib response in sarcoma patients or identifying therapy-selection biomarkers in chordoma were performed on a dataset of limited size. All these aspects are expected to improve as the cohort grows, and as ongoing clinical follow-up studies are published^43^. The actionability of intracellular kinase activity poses a challenge that is often overlooked in single-cohort studies. Because central signaling kinases like ERK, RSK, AKT, and S6K integrate multiple upstream inputs, any one driver contributes only part of the total signal, with the rest reflecting lineage-specific baseline activity and the broader state of an often heavily pretreated tumor. While statistically significant differences in kinase activity are often detectable between genome alteration carriers and non-carriers, these differences do not yet translate into actionable scores at the individual patient level. Realizing this potential will require subtype-aware baselines that go beyond single pan-cancer thresholds toward context-specific refinement and the current study marks an important starting point into this direction. One important area that requires substantial reverse-translational work is to assign function to the thousands of unannotated outlier p-sites detected in patients, and to reduce these to the biomarkers that matter most in a particular cancer context. Systematic phosphoproteome profiling of patient-derived organoids and cell lines under drug or genetic perturbation will likely be instrumental in achieving this, provided it is paired with phenotypic readouts that connect phosphoproteome states to functional outcomes.

Our pan-cancer phosphoproteome resource harbors substantial untapped potential. The immune activity score recapitulated established ICI response patterns across histologies and measures many more parameters than currently employed molecular correlates (MSI, TMB, and PD-L1 mRNA). This warrants retrospective validation of the score in ICI trial cohorts with the potential of improving or even predicting ICI response across cancer sub-types. Another exciting opportunity lies in the cell surface proteome. 94% of patients were positive for at least one targetable protein. Recurrent high expression of established targets (ERBB2, FOLR1, VTCN1, CD276, TACSTD2) outside their approved indications supports repurposing of existing ADC and CAR-T drugs for rare cancers. There are also numerous new proteins with outlier expression that motivates their nomination as targets for preclinical development. With proteome technologies now sufficiently mature, broadly implementing phosphoproteomics in precision oncology is within reach. Integration into controlled basket trials or entity-specific studies is the critical next step to demonstrate clinical significance beyond anecdotal evidence and to validate biomarkers for trial inclusion, particularly for rare cancers where response data are scarce. Shifting advanced molecular diagnostics to earlier treatment lines is another priority because reduced heterogeneity and fewer acquired resistance mechanisms should not only make early-stage phosphoproteomes considerably easier to interpret but also more actionable. In other words, we propose that cancer vulnerabilities, informed by phosphoproteomics as a major descriptor of cancer biology, should be identified and acted upon early in the disease course, prioritizing targeted treatment from the outset rather than as a last resort.

## Resource Availability

## Lead contact

Requests for information/resources should be directed to the lead contact, Bernhard Kuster

## Materials availability

This study did not generate new reagents.

## Data and code availability

All data generated in this study are available via controlled access in the German Genome-Phenome Archive (GHGA).

## Acknowledgements

We thank all members of the TU Munich team, the NCT Sample Processing Laboratory and the DKFZ Next Generation Sequencing, Microarray, and Omics IT and Data Management Core Facilities for high-quality technical and intellectual support and XenoSTART for in vivo PDX studies. This work was supported by the European Research Council (ERC) grant 833710 (TOPAS), the German Federal Ministry of Education and Research (BMBF), grants 01ZZ2322A (PM4Onco), 01KD2207F (HEROES-AYA), 01KD2206C (SATURN-3), 161L0214A (CLINSPECT-M), 03LW0243K (CLINSPECT-M-2), 031L0305A (DROP2AI), 031L0168 (DIAS), 031L0008A (ProteomeTools), the German Science Foundation (DFG) project numbers 325871075 (SFB1309), 329628492 (SFB 1321), 514894665 (TRR387), 452256511 (PromisChemProt), 671907, 641259 (research equipment, grants) and the Chordoma Foundation (grant 3010000958). he MASTER program is supported by the NCT Overarching Clinical Translational Trial Program, the NCT Heidelberg Molecular Precision Oncology Program, and DKTK.

## Author contributions

A.Schneider, S.F., M.T. and B.K. conceived and designed the study. J.W., C.Y.L, S.H., S.W., F.P.B. and N.K. established the (phospho)proteome profiling workflow. J.W., A.Schneider, M.R. and C.S performed and supervised phosphoproteomic sample preparation and LC-MS/MS acquisition. C.J.B., L.D.E., J.B.S., A. Sakhteman, F.H. and M.T. performed (phospho)proteomic data processing and analyses. N.P., L.D., and D.M.F. performed and supervised phenotypic experiments with chordoma cell lines and chordoma PDX models. J.H., B.H., M.O., K.S., C.H., M.Z., M.H. and D.H. analysed WES/WGS and RNA sequencing data. M.V.T., M.W., P.H., S.K., C.E.H., M.H., A.Schneeweiß curated clinical data, discussed all molecular and patient data in MTBs and generated treatment recommendations. K.P., V.T., K.Steiger, C.M. and W.W. established the histopathological workflow and coordinated sample processing. K.B., E.R., V.T., C.S. and A.S. coordinated sample load, prioritization and MTB discussions. C.H.M., A.U., D.J., O.W., U.K., D.R., F.K., A.Stenzinger, S.B., J.T.S., C.B., T.K., E.S., M.B., M.Bitzer, K.S.O., T.H., R.C., R.U., F.L., V.I.G., R.B., V.V., P.W., T.L., V.K., S.H., N.D.; W.J., A. Schneeweiss, P.L., H.G. and S.M.P provided patient tumor material and patient-related metadata. A.Schneider, M.T and B.K. wrote the manuscript. All authors reviewed and edited the manuscript.

## Methods

### 1. Experimental model and study participant details

#### Patient samples

The study included 1998 tumor tissue samples from 1852 patients who provided written informed consent for tumor and control tissue banking, molecular profiling (including germline analysis), and clinical data collection. All procedures were conducted under protocols approved by the Ethics Committee of the Medical Faculty of Heidelberg University (S-206/2011, S-164/2017) and in accordance with the Declaration of Helsinki. Patients were enrolled through one of three programs: the DKFZ/NCT/DKTK Molecularly Aided Stratification for Tumor Eradication Research (MASTER) program, the Comprehensive Assessment of clinical feaTures and biomarkers to identify patients with advanced or metastatic breast Cancer for marker driven trials in Humans (CATCH) program, or the INdividualized therapy FOr high-Risk childhood Malignancies (INFORM) registry. Clinical and molecular data, including demographic information, histopathological and molecular diagnoses, primary tumor and metastasis locations, systemic therapies, staging, molecular tumor board (MTB) recommendations, and application of recommended therapies, were documented in pseudonymized form in a centrally managed electronic data capture system (ONKOSTAR, MASTER program) or a globally accessible web portal (MARVIN, XClinical, INFORM registry).

#### Cell lines

##### Chordoma cell lines

U-CHCF359B (RRID:CVCL_A5IH), CH22 (RRID:CVCL_C159), UM-CHOR5C (RRID:CVCL_A5IC), UM-CHOR5D (RRID:CVCL_A5ID), U-CH2 (RRID:CVCL_4989), 13425-306 (no RRID registered), MUG-CHOR1 (RRID:CVCL_9277), U-CHCF365 (RRID:CVCL_A5II), U-CH17PII (RRID:CVCL_B6R8), MUG-CC1 (RRID:CVCL_AV05), U-CH12 (RRID:CVCL_IR27), U-CH1 (RRID:CVCL_4988), JHC-7 (RRID:CVCL_L154) and UM-CHOR1 (RRID:CVCL_1D68) were used. All cell lines were cultured in IMDM:RPMI-1640 (4:1) supplemented with 10% FBS, 1% NEAA, and 1% GlutaMAX, except for JHC-7 which was cultured in DMEM/F12 supplemented with 10% FBS.

##### Sarcoma cell lines

G-401 (RRID:CVCL_0270), Hs 729.T (RRID:CVCL_0871), KHOS/NP (RRID:CVCL_2546), and RD-ES (RRID:CVCL_2169) were used for reference sample production and obtained from ATCC, CLS (CLS Cell Lines Service GmbH, Germany), and DSMZ (German Collection of Microorganisms and Cell Cultures GmbH, Germany). Cell lines were not independently authenticated for this study but were purchased with authentication certificates from the respective suppliers.

#### Patient-derived xenograft models

Animal experiments were performed at XenoSTART (San Antonio, TX) under IACUC-approved protocols (#09-001, #10-001). Mice were housed at 21.1-23-3°C with 30–60% relative humidity and a 12 h light/dark cycle. Female athymic nude mice (6–12 weeks old; Charles River Laboratories) were implanted subcutaneously with low-passage tumor fragments. Once tumors reached approximately 150–300 mm³, animals were randomized into control and treatment groups (n = 5–7 per group) matched by tumor volume. Study endpoints were defined as the control group reaching a mean tumor volume of 1,500 mm³ or a pre-specified time point. Individual animals with tumors exceeding 2.5 cm³ were removed from the study per IACUC protocol.

### 2. Method details

#### Phosphoproteome profiling

A detailed protocol with each step of the proteomic sample preparation can be found under doi: dx.doi.org/10.17504/protocols.io.14egn9jn6l5d/v1 (private link for review: https://www.protocols.io/private/6257E3B3F98911EFB8F80A58A9FEAC02).

##### Tumor tissue preparation

Tumor tissue was collected following standardized protocols at each contributing site. Material from primary tumors, metastases and locally recurring tumors (material from biopsies and resections) were included. Immediately after collection, samples were snap-frozen without any medium to preserve their integrity.

In the MASTER program, samples collected in Heidelberg (HD) were sent directly to the pathology department via the hospital’s pneumatic tube system. Samples from other contributing sites were shipped to the NCT Sample Processing Lab (SPL) in appropriate containers on dry ice. To evaluate the tumor tissue content, samples were subsequently transferred to the pathology department for embedding in Tissue-Tek® O.C.T.™ embedding medium, placed on top of cork plates and prepared for HE-staining, and then transferred back to the SPL. At the SPL, 50 µm tissue sections were taken until up to 30 mg of tissue was collected. To each sample, a 5 mm metal bead and 800 µl of RLT+ Buffer containing 1% ß-mercaptoethanol (Allprep DNA/RNA/miRNA Universal Kit or QIAamp DNA mini (QIAGEN)) were added. Tumor samples were then mechanically disrupted using a TissueLyser (QIAGEN) (20 Hz, 2 minutes). To remove debris, the lysates were centrifuged (17,000xg, 3 minutes, room temperature) and supernatants were subsequently divided into 600 µl aliquots for DNA and RNA extraction and 200 µl aliquots for phosphoproteome profiling (stored and shipped at −80°C).

In the INFORM program, cryosamples were fixed on cork plates using Tissue-Tek® O.C.T.™ upon arrival and one HE-stained slide was transferred to the neuropathology department to evaluate tumor cell content and necrosis. If tumor cell content was sufficient for analysis (>40%), consecutive slices (10µm) were cut for DNA (20x), RNA (60x), and protein (30x) extraction. Tissue sections were stored at −80°C until shipment for phosphoproteome profiling. For lysis, tissue slices were solubilized in 4% SDS, 40 mM Tris/HCl (pH 7.6), ultrasonicated, and heated (10 min, 95°C). After lysis, the remaining DNA was sheared by adding trifluoracetic acid (TFA) (2 % final concentration, 2 min, 80°C). Next, the lysate was quenched with 4-methylmorpholine (4% final concentration) to neutral pH and the remaining cell debris was removed by centrifugation (20,000 rpm, 20 min).

##### Protein concentration and sample allocation

After lysis, protein concentration was determined by a Bradford (Qiagen Allprep Lysate) or BCA (SDS-lysate) assay, and samples with < 200 µg total protein amount were excluded from further processing. For each TMT-Batch (for TMT see below), 9 patient tumor samples (earlier in the program 8 tumor samples, TMT-channel 1-8/9), which were assigned to batches according to clinical priorities, and two QC reference samples (earlier in the program three reference samples, TMT-channel 9/10-11) were processed in parallel. QC reference samples consisted of 200 µg protein from an equal mix of sarcoma cell lines (RD, HS-729, G-401, KHOS-NP), produced as large stocks (four stocks of sarcoma cell line mixture were produced in this study). One sample of the QC reference was added to the workflow as lysate before protein digestion (TMT-channel 9, batch 1 to 207) and as tryptic peptides at the step of TMT multiplexing (TMT-channel 10/11, all batches).

##### Protein digestion, TMT labeling and multiplexing

Protein digestion was performed using the SP3 approach^87^ and automated on an AssayMAP BRAVO liquid handling platform (Agilent) running a customized protocol. Briefly, 200 µg protein per sample was processed. Proteins in the lysate were precipitated on carboxylate and amine-modified magnetic beads (Sera-Mag A and B, Sigma Aldrich) using 70% ethanol and a 1:5 protein-to-bead ratio. Following reduction and alkylation of cysteins, protein digestion was performed overnight using trypsin (Roche) in a 1:50 (w/w) enzyme-to-substrate ratio. Tryptic peptides were desalted using solid phase extraction (SPE) on Oasis HLB 96 deep well plates (Waters). Next, tryptic peptides of each patient sample were stable isotope labelled using tandem mass tags (TMT) as described^88^. Combined TMT-peptides of patient samples and QC reference samples were desalted by SPE using 50 mg tC18 RP cartridges (Waters).

##### Offline HPLC fractionation and p-peptide enrichment

TMT-labeled samples were separated by high pH reversed-phase HPLC as described^89^ on an X-Bridge BEH130 4.6×250 mm C18 reverse-phase column (Waters) running at a flow rate of 1 mL/min and collecting 96 fractions á 30 seconds that were subsequently pooled into 48 fractions for proteome measurement (14% of the total sample) and 12 fractions (86% of the total sample) for p-peptide enrichment. The latter was accomplished by immobilized metal affinity chromatography using Fe(III)-NTA-IMAC cartridges on the AssayMAP BRAVO liquid handling platform.

##### LC-MS/MS data acquisition

TMT-labelled tryptic peptide batches were analyzed by liquid chromatography tandem mass spectrometry (LC-MS/MS) on Ultimate 3000 or VanquishNeo HPLC systems coupled to Orbitrap Fusion Lumos or Orbitrap Eclipse mass spectrometers (ThermoFisher Scientific). Briefly, for proteome analysis, peptides were separated using a 25-minute solvent gradient and an Acclaim PepMap 100 C18 column running at 50 µL/min^90^. For phosphoproteome analysis, peptides were separated using a 90-minute solvent gradient and a Reprosil Gold column running at 300 nL/min^91^. In both cases, the mass spectrometer was operated in data dependent acquisition (DDA) mode. Intact peptide mass spectra (MS1) were recorded in the Orbitrap. Fragment ion spectra (MS2) were generated by collision induced dissociation (non-modified peptides) or multi-stage activation (phosphopeptides) followed by TMT reporter ion spectra using synchronous precursor selection (MS3).

##### Quality metrics

In the proteome analysis, tumor samples with a summed TMT reporter intensity below 20% of the mean summed intensity of both reference channels were labeled “QC failed” and excluded from the cohort. Full batches were likewise excluded if TMT labeling efficiency was <90% or mean IMAC enrichment efficiency was <80% (calculated as summed phosphopeptide intensity / summed total peptide intensity across 12 fractions). Samples were additionally excluded if contamination was detected in other molecular data layers or if sample identity could not be confirmed (labeled “excluded – other”). Samples with a miscleavage index > 1 (i.e. the median intensity of miscleaved peptides was 10x higher than in the QC channels) were labeled as “miscleavage QC failed”. The miscleavage index was calculated by first correcting log10-transformed peptide intensities from the proteome analysis for relative protein abundance. This correction was performed by subtracting the log10 fold change of the corresponding protein. Then, a miscleavage index was calculated as the median intensity of peptide sequences with at least 1 miscleavage that was quantified in at most 250 patients. Finally, these miscleavage indices were normalized by subtracting the mean miscleavage index of the QC channels in the batch.

Samples with tumor cell content <20% (TCC category “very low”) or a summed TMT reporter intensity below 20% of the mean reference channel intensity in the phosphoproteome analysis were labeled “limited quality.” These samples were included in molecular tumor board (MTB) discussions, with manual corrections applied and heightened interpretive caution, but excluded from cohort-level analyses.

##### TOPAS Pipeline

Data were processed using the TOPAS pipeline as described in the accompanying manuscript (TOPAS: phosphoproteome data analysis and decision support platform for molecular tumor boards). Briefly, raw files were searched with MaxQuant v.1.6.12.0 against the human SwissProt+TrEMBL database (97,057 sequences, downloaded Nov 2020) and common contaminants. Batches with PDX samples were searched against a combined database of the previous mentioned and the mouse SwissProt+TrEMBL database (96,821 sequences, downloaded Jul 2023). Hits mapping to mouse only were removed from evidence.txt file before following analysis. All TMT batches were submitted together to SIMSI-Transfer version 0.5.0^92^ for MS2 identification transfer. Protein and phosphopeptide quantification were performed as described in 3. *Quantification and statistical analysis*. Proteins of interest were annotated as described below. Phosphopeptides were annotated using the Python package psite_annotation v0.5.3 (https://github.com/kusterlab/psite_annotation) using the following files downloaded from PhosphoSitePlus (https://www.phosphosite.org/staticDownloads, accessed January 13th, 2024): Phosphosite_seq.fasta, Kinase_Substrate_Dataset, Phosphorylation_site_dataset, Regulatory_sites.

##### TOPAS scores

Tumor proteome activity (TOPAS) scores were computed as described in the accompanying manuscript (TOPAS: phosphoproteome data analysis and decision support platform for molecular tumor boards). Briefly, the kinase abundance score is the z-score of that patients’ kinase abundance computed relative to the background cohort. The activity indicating phosphorylation score is the sum of log10 fold changes of a set of curated phosphopeptides from literature, drug perturbation experiments and differential expression analysis based on patients in the cohort with an oncogenic RTK alteration, z-scored across all patients. The protein phosphorylation score is the sum of log10 fold changes of all phosphopeptides unique to the protein, z-scored across all patients. The substrate phosphorylation score is the sum of protein-expression-corrected log10 fold changes of all kinase substrates retrieved from *Bayer et al.*^28^, z-scored across all patients. The TOPAS receptor tyrosine kinase score (*TOPAS-RTK*) is the sum of the kinase abundance, activity indicating phosphorylation (weighted 2x) and protein phosphorylation scores. The TOPAS intracellular kinase score (*TOPAS-ICK*) is simply the substrate phosphorylation score. TOPAS scores are again z-scored across all patients to arrive at a standardized TOPAS score for each of the 18 RTKs and 28 ICKs.

##### Immune activity score (IAS)

To quantify the overall immune-related activity of each tumor, we derived a composite Immune Activity Score (IAS) from three biologically defined subscores: antigen presentation machinery, T cell infiltration, and T cell activation. Each subscore was built from a curated feature set spanning three categories (protein abundance, protein phosphorylation score, and substrate phosphorylation score), with individual features assigned a priori to one of the three categories based on established biological function, assembled from Literature (**Table S11**). To compute protein and substrate phosphorylation scores, log2 fold changes relative to the background cohort were summed up and z-scored across the cohort. Also, protein abundances were z-scored across all patients. For each patient, a subscore was then computed as the sum of the z-scored feature values assigned to that category, again z-scored across patients. The composite IAS was calculated per patient as the sum of the three subscores (antigen presentation machinery + T cell infiltration + T cell activation). The resulting per-patient composite values were z-scored once more across all patients, yielding a standardized Immune activity score used in downstream analyses.

#### Genomics and Transcriptomics

##### Sample preparation and sequencing

DNA extraction was performed from fresh frozen tumor tissue using the AllPrep DNA/RNA/miRNA Universal Kit (Qiagen) or QIAamp DNA Mini Kit (Qiagen). In case of formalin-fixed paraffin-embedded (FFPE) tumor samples, DNA was extracted with the GeneRead DNA FFPE Kit (Qiagen). DNA from peripheral blood was extracted using the QIAamp DNA Blood Mini Kit or QIASymphony DSP DNA Mini Kit (Qiagen). DNA quantification and quality control were performed using a Qubit 2.0 Fluorometer (Invitrogen) and a TapeStation 2200 system (Agilent). Whole-genome sequencing (WGS) libraries were prepared from 100 ng genomic DNA using the Illumina TruSeq Nano DNA Library Prep Kit, whereas whole-exome sequencing (WES) libraries were generated from 200 ng DNA using the Agilent SureSelect All Exon v5 or v5 + UTRs Kit. Libraries were sequenced on Illumina HiSeq platforms (HiSeq X Ten, HiSeq 2000, HiSeq 2500, or HiSeq 4000) with paired-end reads. Tumor and matched blood samples were sequenced across two lanes to ensure depth and uniformity. Samples were processed within the NCT Heidelberg Molecular Precision Oncology Program, where the preparation was performed at the Sample Processing Laboratory (SPL), sequencing at the DKFZ Genomics and Proteomics Core Facility (GPCF), and data management was performed through the Omics IT and Data Management Core Facility (ODCF).

##### Data processing and analysis

Logistic and analytic pipelines for molecular profiling in INFORM, MASTER and CATCH have been previously established and described^6,8,93,94^.

In brief: After sequencing DNA, reads were aligned to a reference genome, comprising the human genome (1000 Genomes Phase 2, Genome Reference Consortium; ver. hs37d5) and the Enterobacteria phage phiX174 genome using BWA-MEM (ver. 0.7.15). All parameters were used with default values, except the threshold of minimum score to report alignment (-T) was set as 0. BAM files were sorted with bamsort (biobambam; ver. 0.0.148), and duplicate reads were marked using markdup (Sambamb; ver. 0.6.5^95^). Copy number analysis was performed with CNVkit v2.1.0^96^ for exome sequencing and ACEseq v5.1.0 for whole genome sequencing^96,97^. Segments with a TCN of at least 0.7 above the ploidy were defined as gains (’GAIN’) and segments with a TCN at least 0.3 below the ploidy were defined as losses (’DEL’). Segments with a TCN < 0.5 are called homozygous deletions (’HDEL’), while ‘DEL’ is used for partial copy number losses. A gain is labelled as amplification (‘AMP’), when the TCN is at least 2x base ploidy + 1. Gene fusions (FUS’) were called with the tool Arriba version 0.8^98^. Structural variants (‘SV’) were identified using the in-house (MASTER) developed tool Sophia (https://bitbucket.org/utoprak/sophia/src/master/). Small nucleotide variants(‘SNV’) were called with an in-house pipeline as described in^6^. The number of affected copies was calculated based on the variant allele frequency, copy number at the position of the variant, and the tumor ploidy. If less than 0.5 reference alleles are left, it was assumed that all copies are affected. Homozygous gene deletions and small variants leading to a frameshift, stopgain or aberrant splicing and affecting all copies were interpreted as loss-of-function (LOF). Variants with potential LOF were fusions or structural variants in combination with a deletion or LOH, splicing variants in samples without RNA-Seq, variants of unknown significance, frameshift or stopgain variants if wild-type copies are left. Partial copy number deletions (’DEL’) were not deemed to be sufficient to cause LOF in tumor suppressor genes. Tumor mutational burden (TMB) was defined as the sum of non-silent single-nucleotide variants (SNVs) and coding indels detected per sample. To express TMB in mutations per megabase, this sum was divided by the effective length of the coding genome. Coding regions were defined using GENCODE Human (ver. 19) annotation, and their total length was calculated using bedtools (ver. 2.27.1). For whole-genome sequencing (WGS), the merged coding sequence length was 35.334619 Mb. For whole-exome sequencing (WES), coding regions were additionally intersected with the corresponding target capture design, resulting in coding lengths of 31.057260 Mb for SureSelectXT Human All Exon V5, including UTRs, and 30.894643 Mb for the same design excluding UTRs.

##### OncoKB annotation

OncoKB annotations were retrieved from the OncoKB API v1 (https://api.oncokb.org, accessed August 21st 2025), through the endpoints */api/v1/annotate/mutations/byProteinChange* (SNVs), */api/v1/annotate/copyNumberAlterations* (CNVs), and */api/v1/annotate/structuralVariants* (fusions), passing gene names as *hugoSymbol* parameter.

#### Cell proliferation assays

Chordoma cell lines (U-CHCF359B, CH22, UM-CHOR5C, UM-CHOR5D, U-CH2, 13425-306, MUG-CHOR1, U-CHCF365, U-CH17PII, MUG-CC1, U-CH12, U-CH1, UM-CHOR1) were cultured as described above. For proliferation assay, cells were seeded in a flat-bottom, white-walled, 96-well plate (Corning, Cat# 07-200-566). Assay parameters were optimized for each cell line using Incucyte Live-Cell imaging as follows: the seeding density of each cell line was chosen to give 15% confluence 36 hours after plating, and the assay endpoint was determined as the time at which control cells reach 80%-90% confluence. Afatinib or DMSO vehicle was added 36 hours after seeding, once cells were fully adherent. 10 mM afatinib in DMSO (MedChemExpress; Cat#: HY-10261) was diluted to 3 µM, the top concentration used in this assay, and then serially diluted by 1/3rd to 0.457 nM. At endpoint, cell viability was assessed using CellTiter-Glo (Promega; Cat#: G9242) according to manufacturer instructions, and luminescence was measured using a BioTek Cytation 5 plate reader. For U-CHCF359B cells, which take 20+ days to reach assay endpoint, afatinib resistance was separately confirmed using Incucyte Live-Cell imaging; cells treated with 0.3 µM afatinib displayed minimally inhibited cell proliferation compared to cells treated with DMSO. Data were normalized to DMSO control wells and plotted in GraphPad Prism using log(inhibitor) vs response – variable slope (4 parameter). Absolute EC50 values are reported as the drug concentration required to reduce viability by 50% compared to DMSO-treated control cells. Proliferation assays were repeated at least once for each cell line, for a total of n=2-3 independent biological replicates with consistent results. To yield enough protein for (phospho)proteome profiles, cells were seeded in 10cm² dishes and grown to 80% confluence. Cells were washed twice with PBS and lysed in SDS lysis buffer (2% SDS in 40 mM Tris-HCl, pH 7.6). For DNA hydrolysis, samples were heated at 95 °C for 10 min and incubated with 2% trifluoroacetic acid (TFA) for 1 min, before the reaction was quenched using 4% N-methylmorpholine (NMM). Lysate was cleared by centrifugation at 10,000 × g for 10 min and protein concentration was determined using the Pierce BCA Protein Assay Kit (Thermo Scientific) and cells were further processed as described for patient’s tumor tissue specimen in separate cell line TMT-batches.

#### Xenograft efficacy studies

Animal experiments were performed at XenoSTART in San Antonio, TX in tumor-bearing mice under an institutional animal care and use committee (IACUC)-approved protocol (#09-001, #10-001). Housing conditions for the mice included a room temperature of 70–74 °F, 30–60% relative humidity, and 12 h light/dark cycles. Six to twelve weeks old female athymic nude mice (Charles River Laboratories) were implanted subcutaneously with low-passage tumor fragments. When tumors reached approximately 150-300 mm^3^, animals matched by tumor volume (TV) were randomized into control and treatment groups, each group containing 5-7 animals. Afatinib was purchased from MedChemExpress or LC Labs and formulated in 0.5% methylcellulose, 0.4% Tween 80. Dosing began at Day 0 and afatinib was administered orally at the dose levels and schedules noted in the main text. Animals were observed daily, and weights and TVs were measured twice a week using an electronic scale and digital calipers, respectively. Tumor dimensions were converted to TV using the formula: TV (mm^3^) = width2 (mm) x length (mm) x 0.52. The study endpoints were when the control group reached mean TV = 1500 mm^3^ or a specified time point. Any individual animal reaching a tumor size >2.5 cm^3^ was subsequently removed from the study, in accordance with the XenoSTART IACUC protocol. For mice that reached the tumor volume endpoint before the study endpoint (one vehicle-treated mouse on day 25 of the CF539/TNO155 study), the final TV measurement was carried over and plotted for the remainder of the study, and body weight measurements were not plotted for this mouse beyond this date. To compare treatment efficacy across PDX models with varying growth rates, we used the tumor/control tumor volume (T/C TV) metric, which is discussed in the NCI PDXNet Consensus Recommendations^99^ and is calculated as follows: T/C TV= (T_f_/T_i_)/(C_f_/C_i_), T=mean tumor volume in treatment group, C= mean tumor volume in control group, f=final timepoint (endpoint), i=initial timepoint. Tumor tissue specimens for phosphoproteomic analyses were surgically extracted at study endpoint, flash-frozen in liquid nitrogen and processed as described above for patient material in separate xenograft TMT-batches.

#### Clinical follow up

##### Pazopanib follow-up

Patients for the pazopanib follow-up cohort were pre-selected based on sarcoma diagnosis (oncotree classification) and application of pazopanib with documented start and end of therapy before April 2026 (n=177). The following patient samples were excluded from the cohort (**Table S10**): (1) sample taken under/after pazopanib (n=78), (2) kinase-targeted therapie(s) or multiple lines (>=2) of therapies between sampling and pazopanib start (n=29), (3) 0% Tumor cell content (n=3), (4) incomplete follow-up (n=3), (5) combination therapies (n=2) and (6) KIT-mutant GIST with prior KIT-targeted therapies (n=2). Progression-free survival (PFS) under pazopanib therapy was available for n = 59 sarcoma patients. PFS was defined as the time elapsed from the date of pazopanib therapy initiation to disease progression as defined by local standards and treating physicians. No systematic RECIST evaluation of response was available, and there was no central radiologic review. Patients who discontinued therapy due to treatment-related toxicity (n = 4) or whose reason for discontinuation was unknown (n = 3, loss to follow-up) were censored at the time of therapy discontinuation. A total of 52 events were observed (50 PD, 2 deaths, **Table S10**). Patients were stratified into TOPAS-positive and TOPAS-negative subgroups based on the maximum TOPAS score across the seven pazopanib target kinases (FLT1, FLT4, KDR, KIT, PDGFRA, PDGFRB, and RET), referred to as the ‘Pazopanib target TOPAS’. To evaluate robustness and avoid reliance on a single arbitrary threshold, patient dichotomization was performed independently using three classification criteria: (i) TOPAS ≥ 1.5 versus < 1.5 (n = 18 and n = 41); (ii) TOPAS ≥ 2 versus < 2 (n = 10 and n = 49); and (iii) a data-driven median split at the cohort median TOPAS score of 1.02 (n = 31 and n = 28). Kaplan-Meier survival analysis, log-rank testing, and Cox regression were performed as described in *3. Quantification and statistical analysis*.

##### Immune checkpoint inhibition (ICI) follow-up

The ICI-exploratory cohort comprised 70 patients stratified by Immune Activity Score (IAS, calculated as the sum of TOPAS-IS scores, z-scored across all patients) into a high group (top 35) and a low group (bottom 35). Clinical data were extracted from institutional records and a pre-specified assessment sheet, collected at baseline and at structured follow-up intervals over a minimum of two years, and adjudicated by an independent physician. Patients were selected based on receipt of immune checkpoint inhibitor therapy (monotherapy or combination, n=34) and availability of documented PFS and/or BOR under ICI (n=31). The interval between biomarker sampling and ICI initiation was unrestricted. PFS was defined as the time from ICI start to the first documented disease progression event. Patients without a documented event were censored at the time of therapy discontinuation or last clinical contact, with the reason for censoring recorded for each case (**Table S8**). PFS was expressed in months (1 month = 30.5 days). Patients were dichotomised into IAS-positive or IAS-negative by a pre-specified median split threshold of −0.0035 (median of the full cohort distribution, inclusive of all patients regardless of follow-up availability). IAS-based stratification was performed as described above for TOPAS (pazopanib) stratification. Kaplan-Meier survival analysis, log-rank testing, and Cox regression were performed as described in *3. Quantification and statistical analysis*. Response categories (complete response, CR; partial response, PR; stable disease, SD; progressive disease, PD) were assigned based on the treating physician’s integrated assessment of all available radiographic and clinical information. Best observed response was analyzed as ordinal (PD<SD<PR<CR). The distribution of best observed response (BOR) per IAS group was displayed as a horizontal stacked bar chart, patients with missing BOR were excluded. Response was categorized as (PR, SD, or PD, with proportions (%) shown within segments. Objective response rate (ORR) was defined as the proportion achieving PR and disease control rate (DCR) as the proportion achieving PR or SD.

#### Proteome-based cancer subtype classification

Binary classifiers for tumor entities with ≥10 samples were generated from the pan-cancer cohort using the Python library scikit-learn. Samples with a NOS classification (n=204) and/or low tumor cell content (n=172) were excluded from the training cohort (n=1998 total), yielding the analytical dataset. Remaining samples were stratified by entity size and split into training and test sets: 75/25 for entities with 10–20 samples, and 70/30 for entities with >20 samples. Training set protein abundances were z-scored and the resulting parameters were applied to both the test and NOS sets. Feature selection and model optimization were performed jointly via grid search over two hyperparameters: L1 ratio (0.3, 0.5, 0.7) and inverse regularization strength C (0.1, 1, 10), yielding nine parameter combinations. For each combination, entity-specific protein features were selected using logistic regression with elastic net regularization across 150 cross-validation resampling iterations. The mean and standard deviation of the protein coefficients were used to compute Wald chi-square statistics, and proteins with Benjamini–Hochberg-corrected p < 0.01 were retained. Predictive performance for each parameter combination was then assessed by fitting the selected proteins in a logistic regression model with ridge regularization, evaluated by repeated 3-fold cross-validation (35 repetitions), and scored by the Matthews correlation coefficient (MCC). The optimal model was selected as the combination achieving the highest training MCC, with preference for fewer proteins within a 0.15 MCC tolerance. Classification thresholds were set by maximizing the F0.5-score on the test set to prioritize precision, then applied to classify test samples. This process was repeated for every tumor entity. Models achieving MCC ≥ 0.7 were subsequently applied to NOS samples; positive classifications were reviewed against orthogonal metadata to assess concordance with other diagnostic modalities.

#### UMAP analysis

Intensities of proteins quantified in >70% of patients were log_10_ transformed. Missing values were imputed using the PPCA method included in scikit-learn v1.3.0. UMAPs were calculated using the Python package umap-learn v0.5.3 using Euclidean distance, 10 nearest neighbors and 1000 epochs.

#### Reproducibility of phosphoproteome analysis

For all proteins or phosphopeptides found in >70% of samples, non-log-transformed intensities were used to compute the coefficient of variation (CV) as a metric for quantitative precision. Cumulative density plots for CV were created for each QC channel across batches for each of the four lots of QC samples (lot 1 for batch 1-72 (+ 117-120), lot 2 for batch 76-179 (excl. 117-120), lot 3 for batch 184-311, lot 4 for batch 315-336).

#### Protein of interest annotation: Data sources

Cancer gene annotations were downloaded from OncoKB (https://www.oncokb.org/cancer-genes, accessed 22 Aug 2024). Kinase gene annotations were downloaded from KinHub (http://www.kinhub.org/kinases.html, accessed 17 Jan 2024). ADC targets were downloaded from ADCdb (http://adcdb.idrblab.net/, accessed 19 Aug 2024). Immunotherapy targets were collected from literature^100–102^. Surface-annotated genes were downloaded from the Wollscheid lab surfaceome resource (SURFY: https://wlab.ethz.ch/surfaceome/),; CSPA: https://wlab.ethz.ch/CSPA/, CSPA validated surfaceome proteins, accessed 27 May 2025) or the Human Protein Atlas resource (https://www.proteinatlas.org/about/download, subcellular_location.tsv filtered to plasma mebrane, accessed 27 May 2025).

### 3. Quantification and statistical analysis

#### Protein and phosphopeptide quantification

TMT-based MS3 reporter intensity quantification was performed by the Python package SIMSI-Transfer v0.6.3 using isotope impurity correction factors provided by the vendor. Within each batch, MS3 intensities were median centered and MS1 intensities were median centered across batches. MS3 intensities per channel were transformed into quasi-LFQ^103^ intensities as (MS1 intensity)*(MS3 intensity in channel)/(summed MS3 intensity across channels). Protein grouping, FDR control and MaxLFQ quantification on all TMT batches together was performed on gene level using the picked protein group FDR package v0.6.2^104^. For phosphopeptides, precursor intensities were summed up across charge states and fractions. Phosphopeptides with the same naked sequence and number of phosphorylated residues were grouped if phosphorylated residues were within a ±2 residue window. For simplicity, these groups are referred to as phosphopeptides throughout the manuscript and the remainder of the methods section. For each protein and phosphopeptide, intensities were log10-transformed and the rank, z-score, and fold change were calculated relative to the cohort.

#### Survival analysis

For the pazopanib and immune activity score analyses, Kaplan-Meier survival functions were estimated for each subgroup using the product-limit estimator. Pointwise 95% confidence intervals were computed using Greenwood’s variance formula. Censored observations are indicated on survival curves as vertical tick marks. Survival distributions between positive and negative groups were compared using the two-sided log-rank test. Hazard ratios (HR) with 95% confidence intervals were derived from a univariable Cox proportional hazards regression model, with the positive group as the test variable and the negative group as the reference category, an HR < 1 therefore indicates a lower instantaneous hazard of progression or death in the positive group. Numbers of patients at risk are reported at 5-month intervals below each curve. All survival analyses were carried out in Python (version 3.13) using the lifelines package^105^. A p value < 0.05 was considered statistically significant.

#### Immune activity score (IAS) statistical analysis

Statistical tests were performed using the Python package SciPy v1.16.2. The Wilcoxon rank-sum test (scipy.stats.mannwhitneyu; two-sided) was applied to numerically coded BOR (PD = 0, SD = 1, PR = 2), with effect size quantified by Cliff’s delta (δ; rank-biserial correlation r = (U₁ − U₂) / (n₁ × n₂)) and bias-corrected and accelerated (BCa) bootstrap 95% CIs derived from 2,000 resamples (with a fixed random seed of 42)A cumulative-link proportional-odds model (CLM; statsmodels OrderedModel, logit link) was fitted with IAS group (IA-positive = 1) as the sole predictor; the odds ratio (OR) with 95% BCa bootstrap CI (B = 2,000) is reported. Group differences between objective response rate (ORR) and disease control rate (DCR) were assessed by a two-sided Fisher’s exact test.

#### Correlation analysis

For continuous, normally distributed outcomes (e.g., expression values), Pearson correlation coefficients (*r*) were calculated to quantify the linear relationship between variables. Statistical significance was determined by a two-tailed *t*-test on the correlation coefficient. For time-to-event outcomes subject to right-censoring (e.g., progression-free survival, PFS), associations were assessed using univariate Cox proportional hazards (CoxPH) models, with the molecular score entered as a continuous covariate. The resulting hazard ratio (HR) and 95% confidence interval (CI) quantify the change in instantaneous risk of the event per unit increase in the score; HR < 1 indicates that higher scores are associated with longer survival. Statistical significance was assessed by the Wald test and confirmed by the likelihood-ratio test. Model discrimination was reported as the concordance index (C-index). All analyses were performed in Python using scipy (v1.9, Pearson correlation) and lifelines (v0.27, Cox model).

#### Differential protein abundance analysis

For each protein, a p-value and fold-change was calculated between the protein abundances of two patient groups (responder and non-responder) for high and low EGFR TOPAS score using the independent t-test function *from the Python package Scipy* v1.16.2. Plots of log2 fold change versus -log10 adjusted *p*-value are shown with annotation of for significantly differentially expressed proteins (p<0.01) with the highest fold-change. From those, TUSC3 and MX1 were selected for treatment stratification of patient samples and models.

---

**Figure S1:**
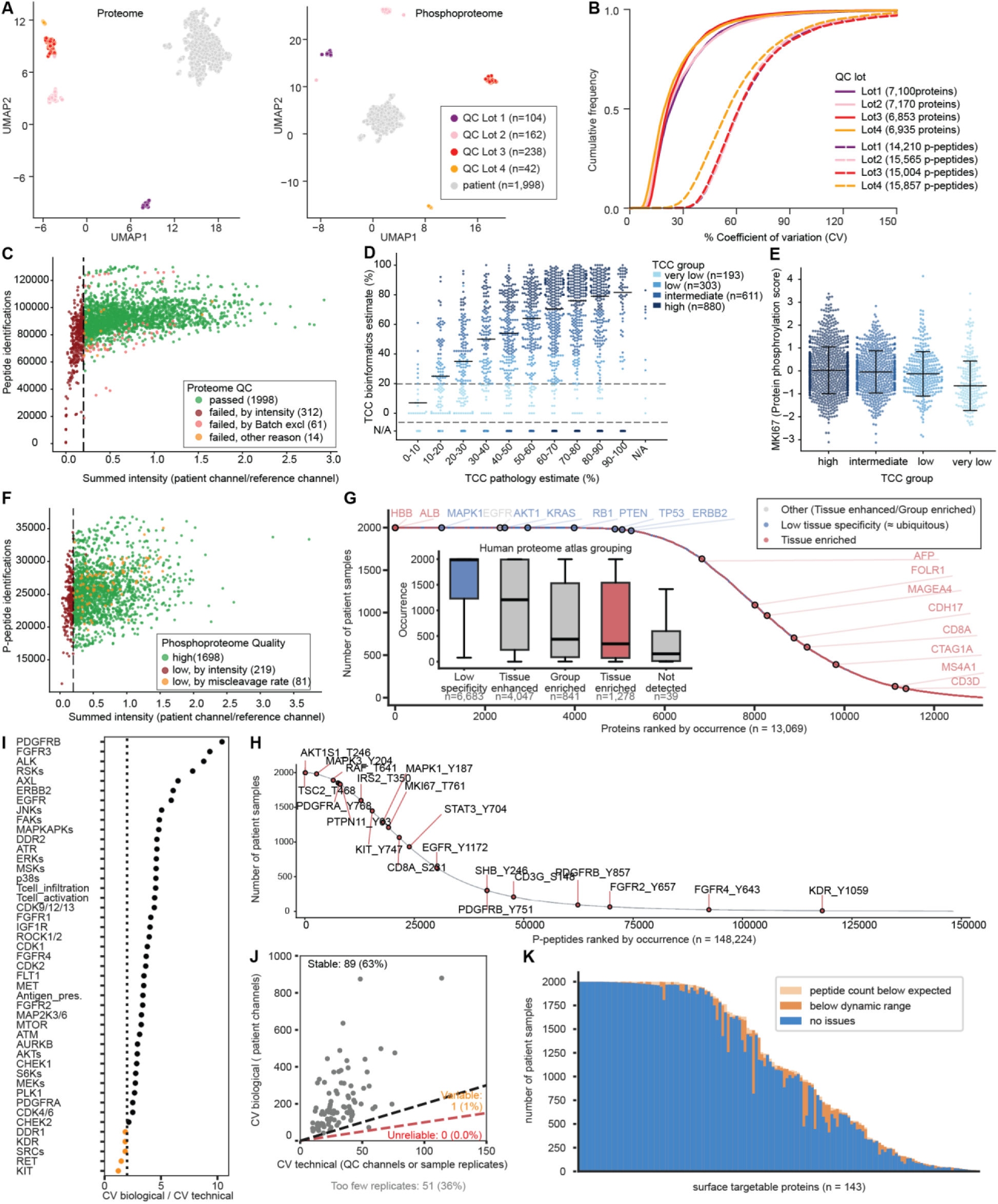
Feasibility of (phospho)proteome profiling in precision oncology trials. (A) UMAP projection of expression data from proteins (left, 70% occurrence across reference channels) and p-peptides (right, 70% occurrence across reference channels), showing clustering of patient tumor samples (grey, n = 1,998) and QC samples from 4 QC lots (colored, n = 42–238). (B) Cumulative frequency plots showing the quantitative precision (coefficient of variation, CV) of technical replicates across 4 QC lots (n = 42–238) at the protein (solid line) and p-peptide (dashed line) level. (C) Scatter plot showing the relationship between peptide identifications (proteome) and summed intensity ratio (patient/reference channel), colored by QC category (passed vs. failed, for different QC criteria). The QC-cutoff ratio of 0.2 is indicated by a dashed vertical line. (D) Relationship between the TCC pathology estimate (x-axis) and the TCC bioinformatics estimate (y-axis), with resulting TCC grouping indicated by color. (E) MKI67 phosphorylation score across TCC groups. Median and standard deviation (SD) are indicated in the plot. (F) Scatter plot showing the relationship between p-peptide identifications and summed intensity ratio (patient/reference channel), colored by QC category (passed vs. failed, for different QC criteria). The QC-cutoff ratio of 0.2 is indicated by a dashed vertical line. (G) Ranked list of the 13,069 quantified proteins in this study (identified at 1% FDR), ordered by decreasing occurrence in patient samples. Selected proteins highlighted in the plot are colored by Human Protein Atlas (HPA) category of tissue specificity: red = tissue enriched, blue = low tissue specificity, grey = other (tissue enhanced, group enriched or not detected in HPA). Inset: box plot indicating the distribution and median of occurrence of proteins across HPA categories. (H) Ranked list of the 148,224 quantified p-peptide groups in this study (identified at 1% FDR), ordered by decreasing occurrence in patient samples. Several p-peptides of interest are highlighted in the plot. (I) Technical variation assessed as the ratio of biological CV (CV across patient samples) to technical CV (CV across QC samples and patient replicates) for 42 TOPAS scores and 3 immune activity subscores. Scores are sorted from highest to lowest ratio; a reliability threshold of 2 is indicated by a dashed vertical line, and scores not passing this threshold are colored in orange. (J) Grouping of surface-targetable proteins of interest (POIs) by quantification robustness as stable (grey), variable (orange), or unreliable (red). Robustness was assessed via biological vs. technical CV for 90 proteins (detected in reference channels or patient replicates). POIs detected in too few technical replicates are not shown in the plot and are listed below it. (K) Ranked list of surface-targetable POIs, ordered by decreasing occurrence in patient samples. POI–patient pairs with no reported issue (blue fraction) are considered for MTB reporting. POI–patient pairs with potential identification issues (light orange; peptide count below expected) or quantification issues (orange;intensity below dynamic range) are flagged in the patient report.

**Figure S2:**
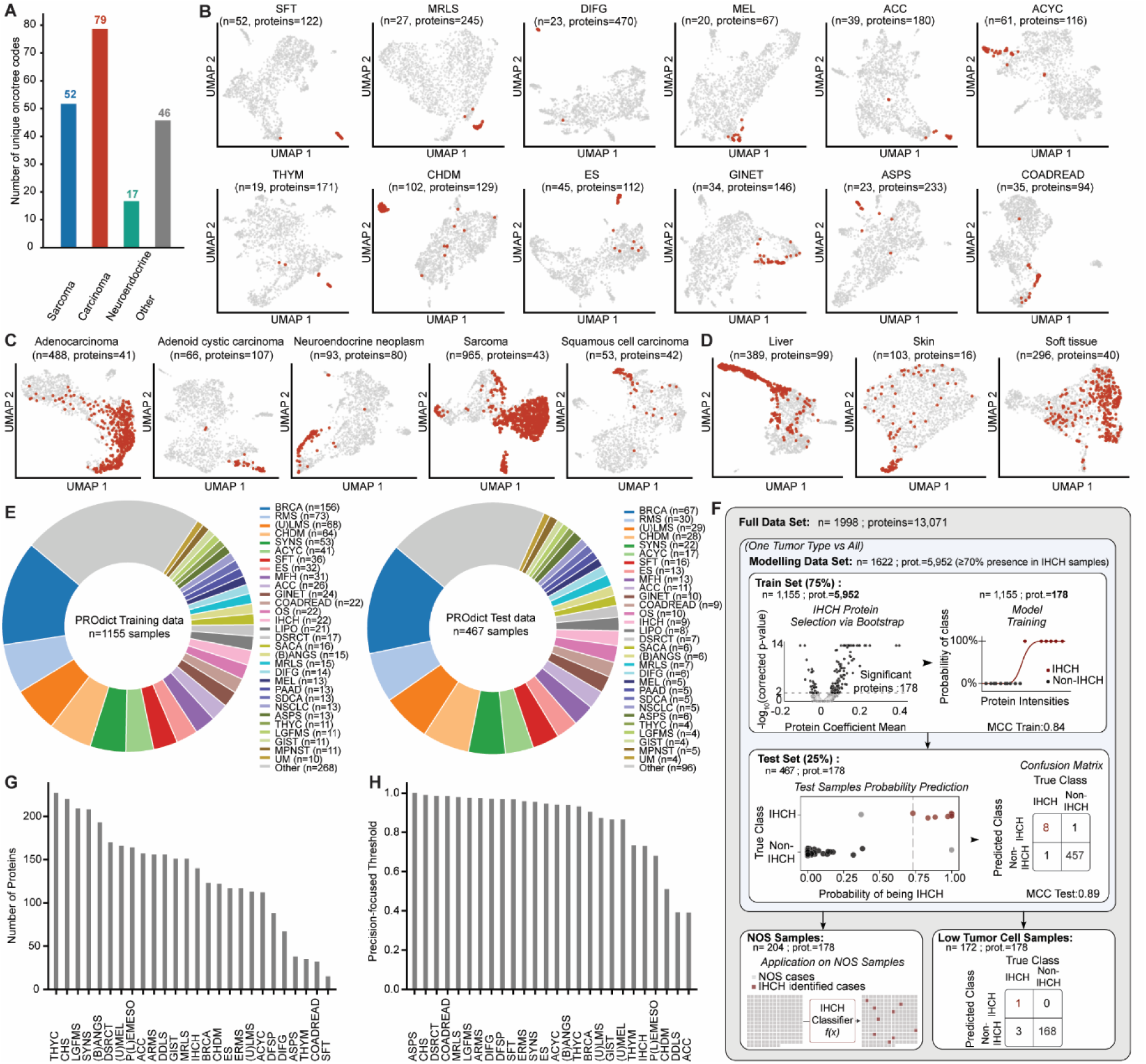
Tumor-of-origin dominates the real-world pan-cancer proteome. (A) Bar plot indicating the number of distinct cancer subtypes (unique OncoTree codes) per broad histologic cancer type: sarcoma, carcinoma, neuroendocrine neoplasms, and other (including melanoma, mesothelioma, germ cell tumors, and thymic epithelial tumors). (B-D) UMAP projections of protein expression data using protein signatures selected per tumor subtype (B), broad histologic subtype (C), or tissue topology (D), each optimized for UMAP separation. The respective tumor subtype/histology/tissue topology is colored red, while all other samples are colored grey. The number of samples and proteins is indicated above each UMAP. (E) Pie charts indicating the size and tumor-subtype composition of the PROdict training (left) and test (right) cohorts. (F) Schematic of the PROdict workflow for intrahepatic cholangiocarcinoma (IHCH) model: protein selection (logistic regression with elastic-net regularization) followed by final model training (ridge-regularized logistic regression), performance evaluation using the test cohort (middle box), and application to re-classification of NOS samples or classification of samples with low tumor cell content (lower boxes). (G) Bar plot ranking protein signature size across all retained classifiers (range: 15–227 proteins). (H) Bar plot ranking the precision-focused classification probability threshold for each retained classifier.

**Figure S3:**
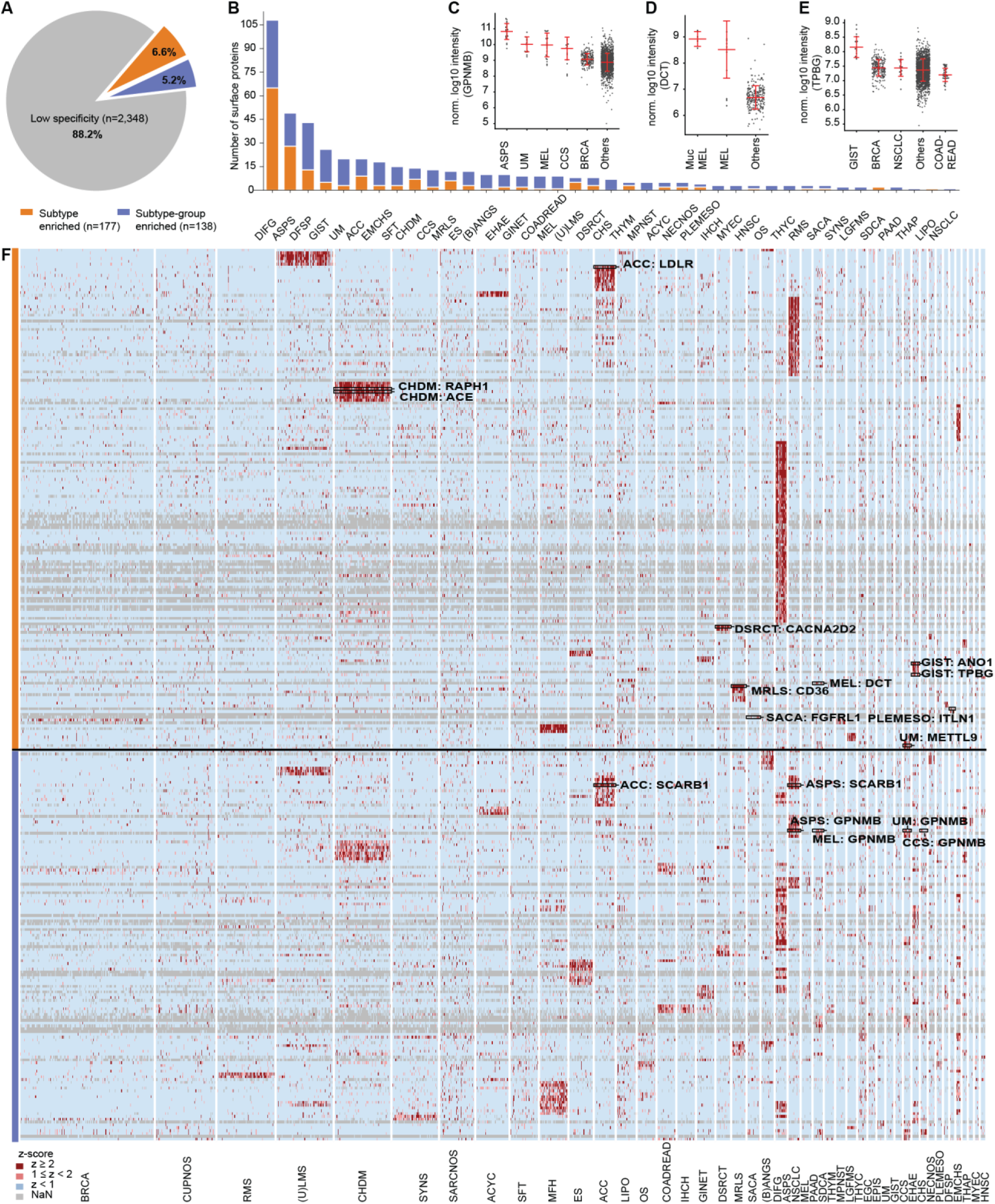
Pan-cancer surfaceome. (A) Pie chart depicting the classification of surface proteins across 45 cancer subtypes into: subtype enriched (median z ≥ 2 in 1 subtype and median z < 1.5 in all 44 other subtypes; orange), subtype group enriched (median z ≥ 1.5 in 2–5 subtypes and median z < 1.5 in all others; blue), and low specificity (expressed broadly across subtypes; grey). (B) Stacked bar chart showing the number of subtype enriched (orange) and subtype-group enriched (blue) surface proteins per subtype, for the 39 of 45 subtypes with n ≥ 1 protein in either category. (C) Normalized log10 intensity of GPNMB (subtype group enriched), comparing the four enriched subtypes (ASPS, UM, MEL, CCS) against BRCA and all others. Median and SD are depicted in the plot. (D) Normalized log10 intensity of DCT (subtype enriched), comparing MEL and mucosal MEL against all others. Median and SD are depicted in the plot. (E) Normalized log10 intensity of TPBG (subtype enriched), comparing GIST against BRCA, NSCLC, COADREAD, and all others. Median and SD are depicted in the plot. (F) Heatmap depicting all subtype enriched (n = 177) and subtype-group enriched (n = 138) proteins across 45 cancer subtypes. Rectangles highlight protein–cancer subtype pairs of interest.

**Figure S4:**
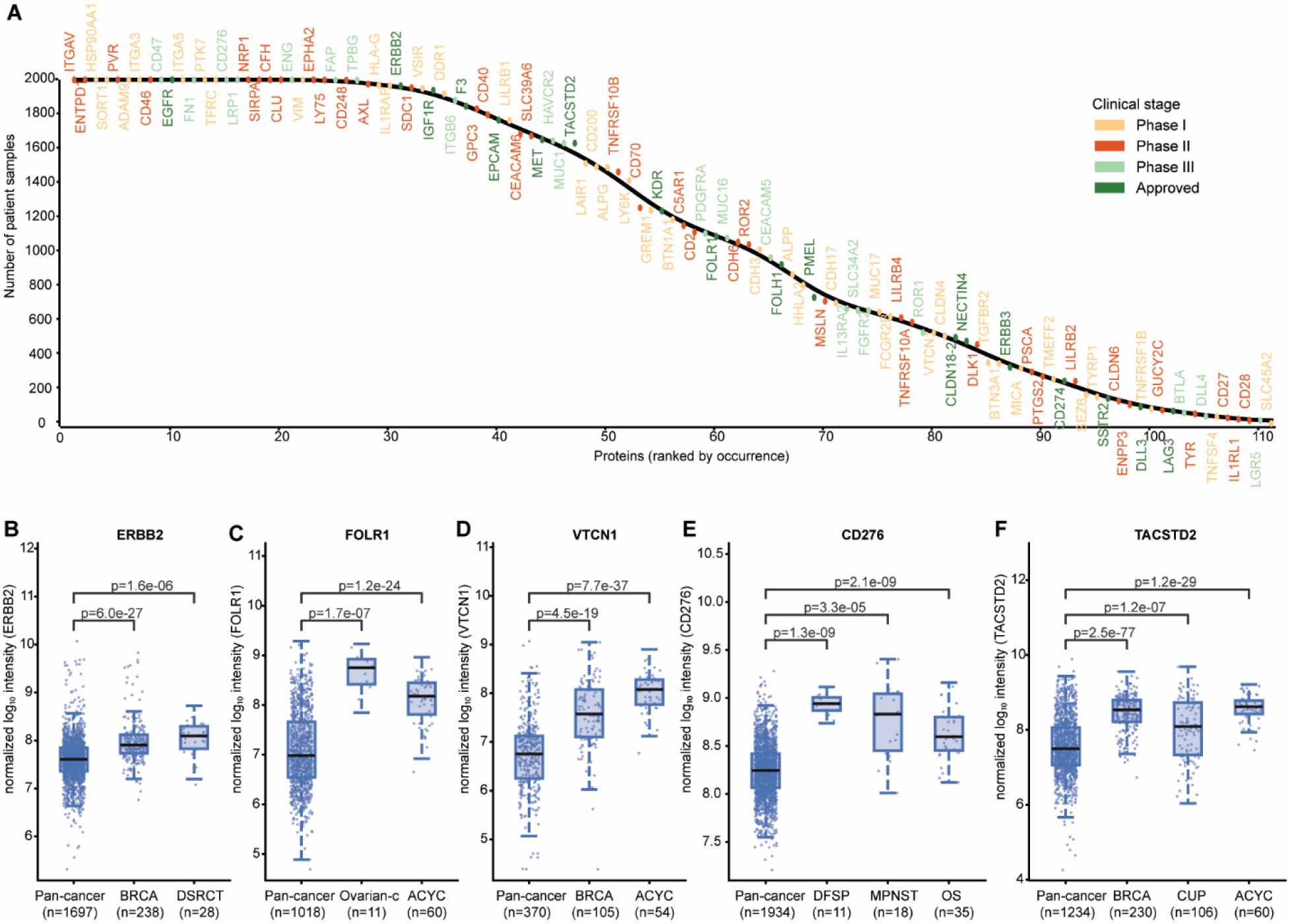
The druggable pan-cancer surfaceome. (A) Ranked list of the 111 surface-targetable proteins quantified in this study (identified at 1% FDR), ordered by decreasing occurrence in patient samples and colored by clinical stage (Phase I–III, approved). (B-F) Box plots depicting the normalized log10 intensity of ERBB2 in BRCA and DSRCT compared to all others (B), FOLR1 in ovarian carcinoma (OncoTree-codes: SOC, EOV, OOVC, OGCT, GRCT, OMGCT, SCCO) and ACYC compared to all others (C), VTCN1 in BRCA and ACYC compared to all others (D), CD276 in DFSP, MPNST, and OS compared to all others (E), and TACSTD2 in BRCA, CUP, and ACYC compared to all others (F). P-values indicating significant enrichment of the illustrated subtypes relative to the pan-cancer cohort are shown in each plot.

**Figure S5:**
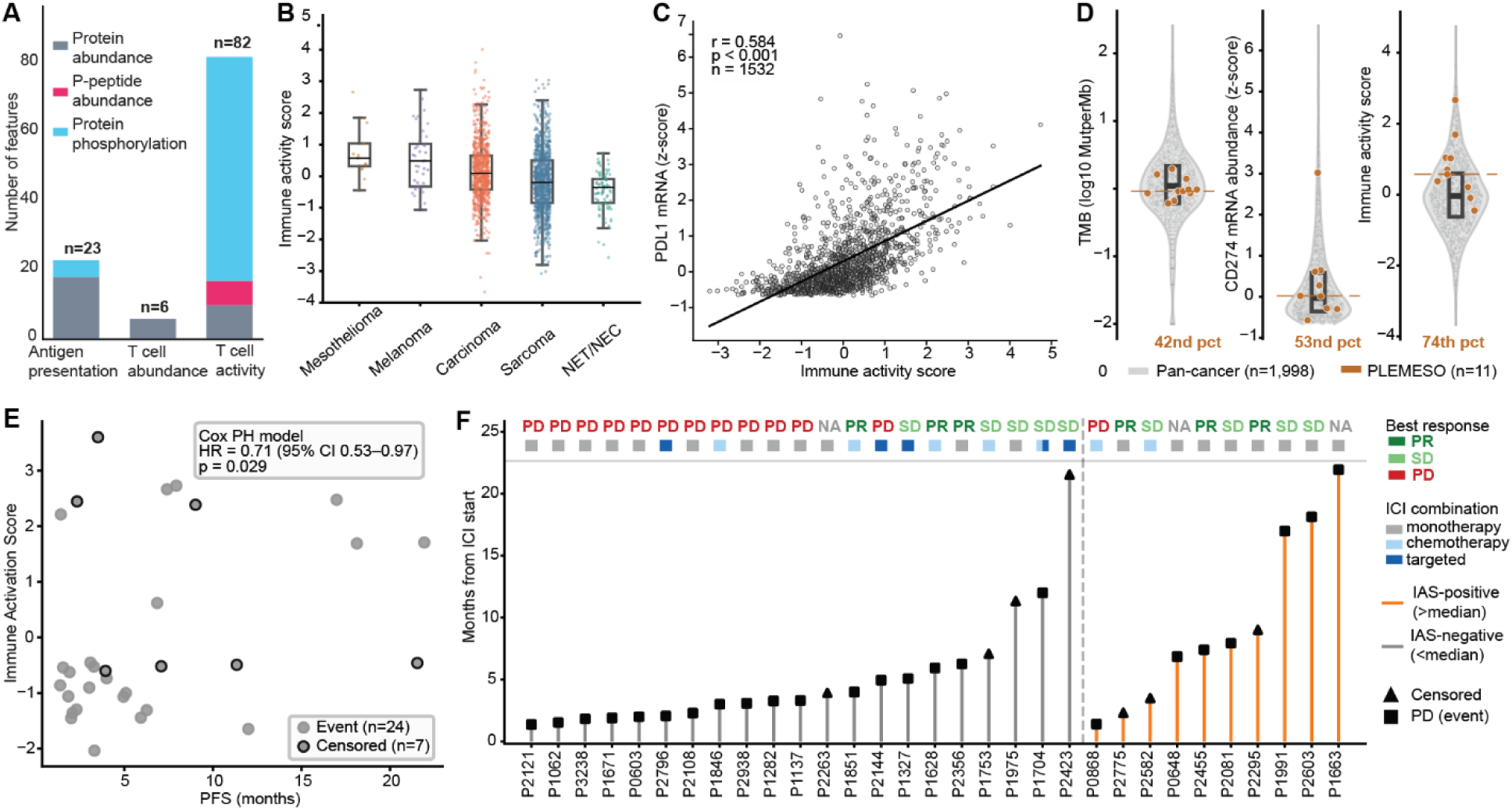
A phosphoproteomics-derived immune activity score captures functional immune states and associates with ICI response. (A) Fraction of protein abundance, p-peptide abundance and phospho-proteins (protein phosphorylation score) scored per immune activity subscore. (B) Immune activity score across a broad range of cancer histologies. Sorted by decreasing median score. (C) Correlation of immune activity score and PDL1 (CD274) mRNA abundance (z-scored FPKM) across 1532 tumor specimens with available transcriptomic data. The Pearson correlation coefficient (r) and corresponding p-value are indicated in the plot. (D) Comparing biomarkers for ICI stratification across the pan-cancer cohort (n = 1,998; grey) with pleural mesothelioma (PLEMESO; n = 11; brown) highlighted. Each panel shows a sina plot in which individual samples are jittered within the local density of the violin. The black box and crossbar indicate the pan-cancer interquartile range and median, respectively. The dashed brown line marks the PLEMESO subtype median, with its percentile rank within the pan-cancer distribution indicated. (E) IAS correlates with PFS under ICI therapy: Scatter plot of immune activation score versus PFS under ICI therapy. Each grey dot represents one patient,dots with black stroke represent censored events. (F) Individual patient progression-free survival under ICI therapy: Swimmer plot showing PFS ICI (months from ICI start) for each patient (n = 31; one patient excluded due to missing PFS data). Patients are grouped by IAS status (IAS-positive: orange lines; IAS-negative: grey lines) and sorted by PFS duration within each group. End-of-follow-up markers: square = PD, triangle = censored. Color bars above each patient line indicate ICI combination type (grey = monotherapy; light blue = chemotherapy combination; dark blue = other combination; split bar = chemotherapy + other). Best observed response is annotated above each color bar (PR: dark green; SD: light green; PD: red; NA: grey).

**Figure S6:**
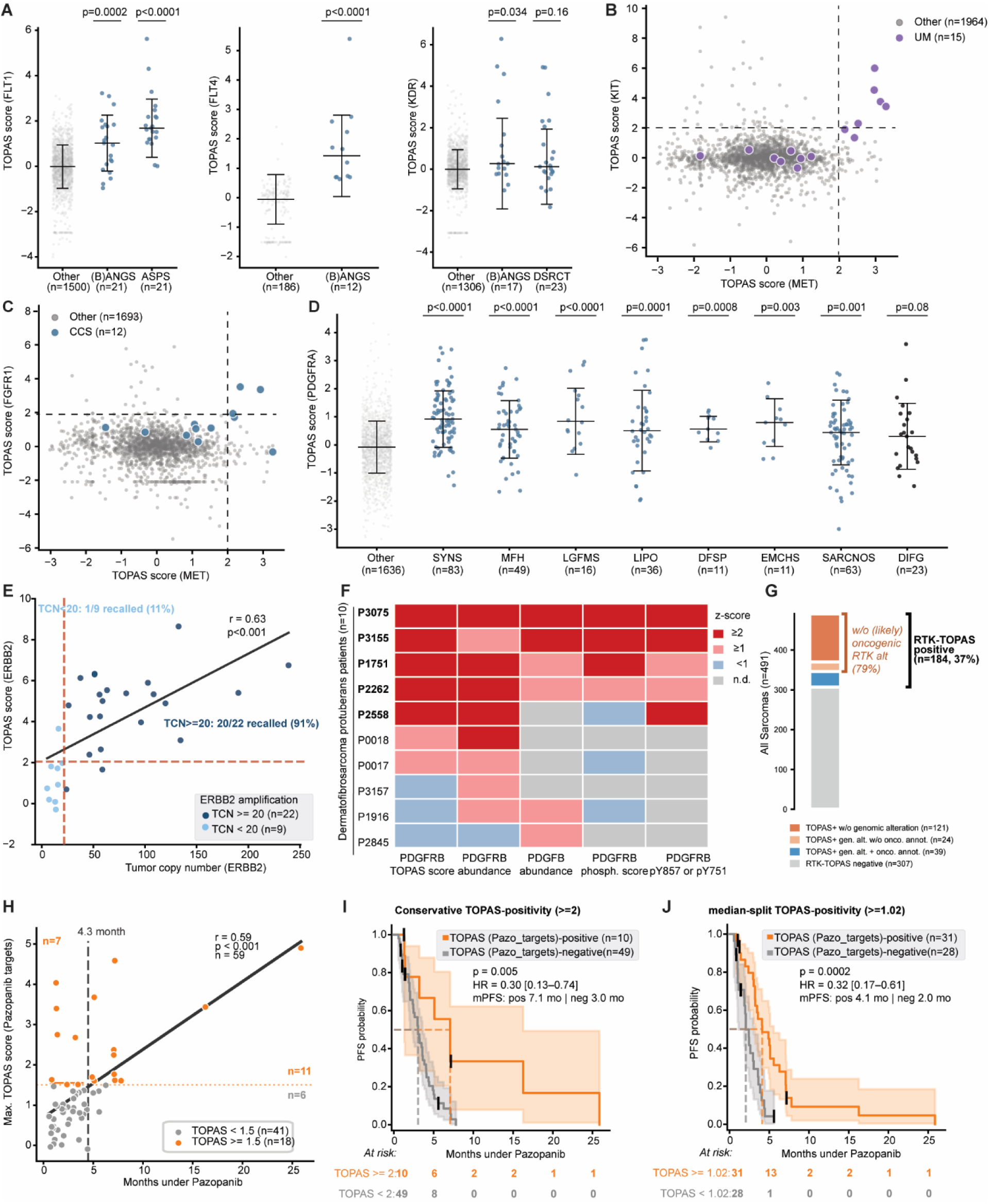
RTK activity and genomic status in advanced cancers. (A) FLT1 (VEGFR1), KDR (VEGFR2) and FLT4 (VEGFR3) TOPAS scores across selected cancer subtypes. Each panel shows kinase scores for one receptor; each dot represents one patient sample. Pan-cancer (Other) is shown in grey; selected sarcoma subtypes are shown in blue. Black horizontal bars show the median; vertical error bars indicate ±1 standard deviation (SD). P-values are from two-sided Mann-Whitney U tests comparing each subtype against pan-cancer (Other). (B, C) RTK co-activation in uveal melanoma (UM,E) and clear cell sarcoma (CCS,F). Scatter plots of pairwise TOPAS scores for individual patient samples. Pan-cancer background is shown in grey; UM in purple and CCS in blue. Dashed lines indicate positivity cut-off for TOPAS scores. (D) PDGFRA kinase activity scores across selected cancer subtypes. Pan-cancer (Other) is shown in grey, Sarcoma subtypes are shown in blue and glioma (DIFG) are shown in black. Black horizontal bars show the median; vertical error bars indicate ±1 standard deviation (SD). P-values are from two-sided Mann-Whitney U tests comparing each subtype against pan-cancer (Other). (E) Scatter plot with linear regression fit depicting the correlation between ERBB2 gene tumor copy number (TCN) and TOPAS score, colored by TCN < 20 (light blue) and TCN ≥ 20 (blue). The dashed line indicates the conservative cut-off for TOPAS-positivity (≥ 2). The Pearson correlation coefficient (r) and corresponding p-value are indicated in the plot. (F) Heatmap showing PDGFRB TOPAS score, PDGFRB and PDGFB protein abundance, PDGFRB protein phosphorylation score, and PDGFRB pY857 or pY751 (max z-score) abundance across 10 patients with dermatofibrosarcoma protuberans (DFSP, PDGFB:COL1A1 fusion). Patients are ranked by PDGFRB TOPAS score (descending from top to bottom). (G) Stacked bar plot indicating the fraction of TOPAS-positive sarcoma patients without a genomic RTK alteration (orange), with an OncoKB-annotated genomic RTK alteration (light orange), and with a genomic RTK alteration not annotated in OncoKB (blue). TOPAS-negative (for all RTKs) sarcoma patients are indicated in grey. (H) Scatter plot depicting the relationship between the maximum TOPAS score across pazopanib target kinases (FLT1, FLT4, KDR, KIT, PDGFRA, PDGFRB, and RET; y-axis) and the duration of pazopanib therapy as a surrogate for progression-free survival (PFS; x-axis) in n = 59 sarcoma patients. Each dot represents one patient. Patients with a TOPAS score ≥ 1.5, classified as TOPAS-positive, are shown in orange (n = 18); patients with a TOPAS score < 1.5 are shown in grey (n = 41). The solid regression line indicates the linear fit across all patients. The dashed vertical line marks a PFS threshold of 4.3 months (considered clinical benefit). (I,J) Kaplan–Meier curves depicting progression-free survival (PFS) probability under pazopanib therapy in a cohort of advanced sarcoma patients (n = 59), stratified by the maximum TOPAS score across pazopanib target kinases (FLT1, FLT4, KDR, KIT, PDGFRA, PDGFRB, and RET). Patients were dichotomized into TOPAS-positive and TOPAS-negative groups using a threshold of ≥ 2 vs. < 2 (e, n = 10/49) or ≥ median (1.02) vs. < 1.02 (f, n = 31/28). PFS was defined as the time from therapy initiation to progressive disease (PD) or death. Patients discontinuing due to toxicity (n = 4) or for unknown reasons (n = 3) were censored and are indicated by vertical tick marks (|). Shaded areas = 95% confidence intervals (Greenwood’s formula). Dashed lines = median PFS. Numbers at risk are shown below the plot at 5-month intervals. HR with 95% CI derived from a univariable Cox proportional hazards model, using the TOPAS-negative group as reference.

**Figure S7:**
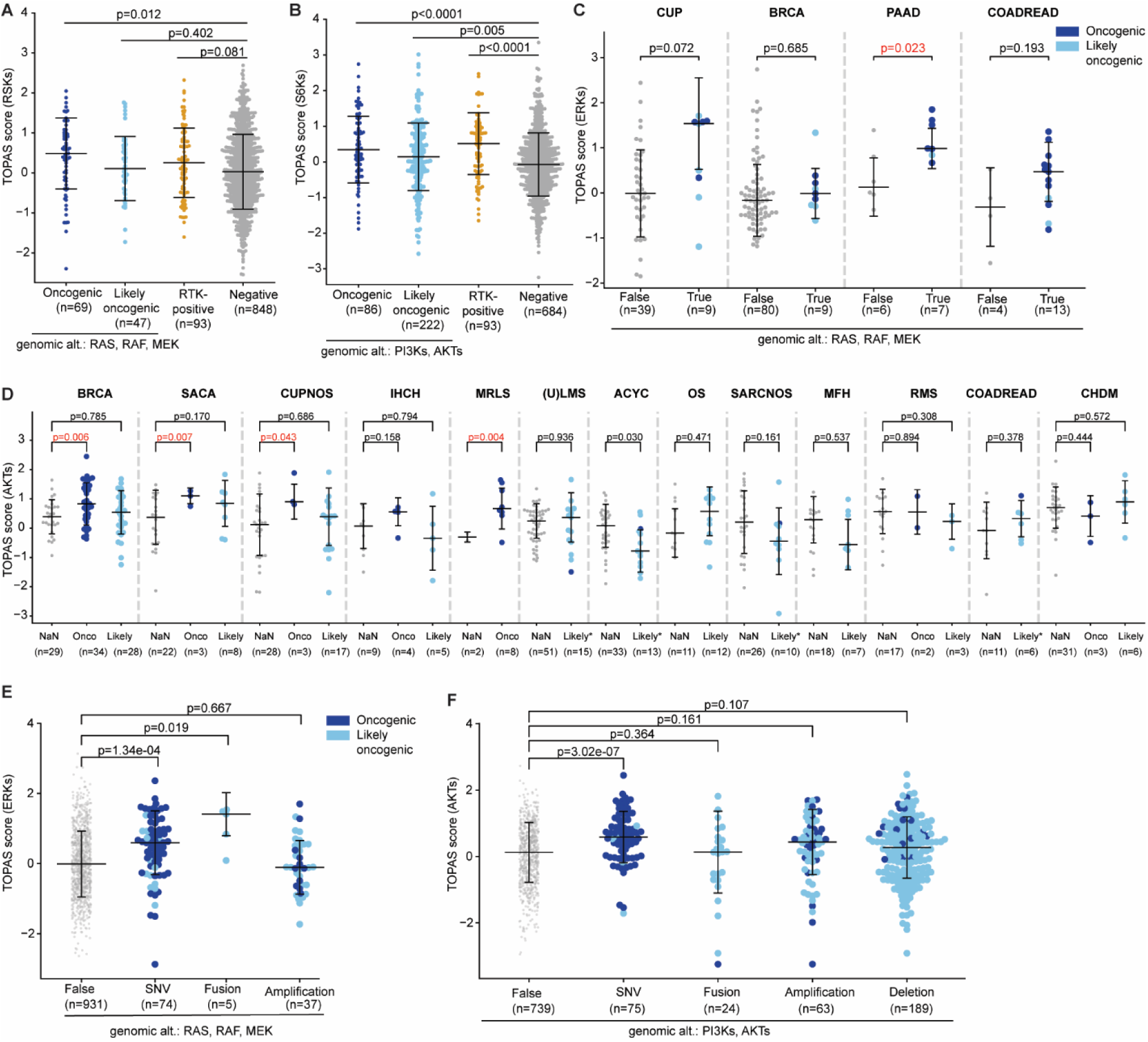
Subtype-resolved landscape of intracellular growth kinase activities. (A) RSKsTOPAS score shown for patient samples grouped by genomic alteration status in RAS/RAF/MEK or RTK-positivity: at least one oncogenic (blue) or likely oncogenic (light blue) alteration in one of the RAS/RAF/MEK isoforms, or RTK-positive (RTK TOPAS score ≥ 2 and a genomic RTK alteration; red), vs. negative (none of the three criteria; grey). Each dot represents one tumor sample. The horizontal bar indicates the median; error bars denote ±1 standard deviation. Statistical comparisons between each group and the negative reference group were performed using a two-sided Welch’s t-test; p-values are shown above brackets. (B) S6Ks TOPAS score shown for patient samples grouped by genomic alteration status in PI3K/AKT or RTK-positivity: at least one oncogenic (blue) or likely oncogenic (light blue) alteration in one of the PI3K or AKT isoforms, or RTK-positive (RTK TOPAS score ≥ 2 and a genomic RTK alteration; red), vs. negative (none of the three criteria; grey). Each dot represents one tumor sample. The horizontal bar indicates the median; error bars denote ±1 standard deviation. Statistical comparisons between each group and the negative reference group were performed using a two-sided Welch’s t-test; p-values are shown above brackets. (C) ERKs TOPAS scores across four cancer subtypes frequently altered in RAS/RAF/MEK (CUP, BRCA, PAAD, COADREAD). Per subtype, samples are divided into those without a RAS/RAF/MEK alteration (No event; grey) and those with an oncogenic or likely oncogenic alteration (oncogenic = blue, likely oncogenic = light blue). Each dot represents one tumor sample. Horizontal bars indicate the median; error bars denote ±1 standard deviation. Statistical comparisons between unaltered (No event) and altered groups were performed using a two-sided Welch’s t-test; p-values are shown above brackets. Only cancer subtypes with ≥ 5 samples carrying an alteration are displayed. Oncogenic alterations were defined according to OncoKB criteria. (D) AKTs TOPAS scores across 13 cancer subtypes with PI3K/AKT pathway alterations. Per subtype, samples are grouped by alteration status: no alteration (grey), oncogenic (blue), and likely oncogenic (light blue). For subtypes with fewer than two oncogenic samples, the oncogenic and likely oncogenic samples were merged into a single group (indicated by Likely*), retaining individual point colors. Each dot represents one tumor sample. Horizontal bars indicate the median; error bars denote ±1 standard deviation. Statistical comparisons between unaltered (No event) and altered groups were performed using a two-sided Welch’s t-test; p-values are shown above brackets. Only cancer subtypes with ≥ 5 samples carrying an alteration are displayed. Oncogenic alterations were defined according to OncoKB criteria. (E,F) ERKs (E) and AKTs (F) TOPAS scores are shown for all tumor samples, grouped by the type of genomic alteration. Samples without any genomic alteration in the indicated isoforms are displayed as a strip plot (No event; grey). Samples harboring single nucleotide variants (SNV), gene fusions (Fusion), or gene amplifications (Amplification) are shown as swarm plots, with individual points colored by oncogenicity classification: oncogenic (blue) or likely oncogenic (light blue), as defined by OncoKB. The horizontal bar indicates the median; error bars denote ±1 standard deviation. Statistical comparisons between the no-event group and each alteration type were performed using a two-sided Welch’s t-test; p-values are indicated above brackets. Samples carrying alterations of multiple event types are counted in each respective group.

**Figure S8:**
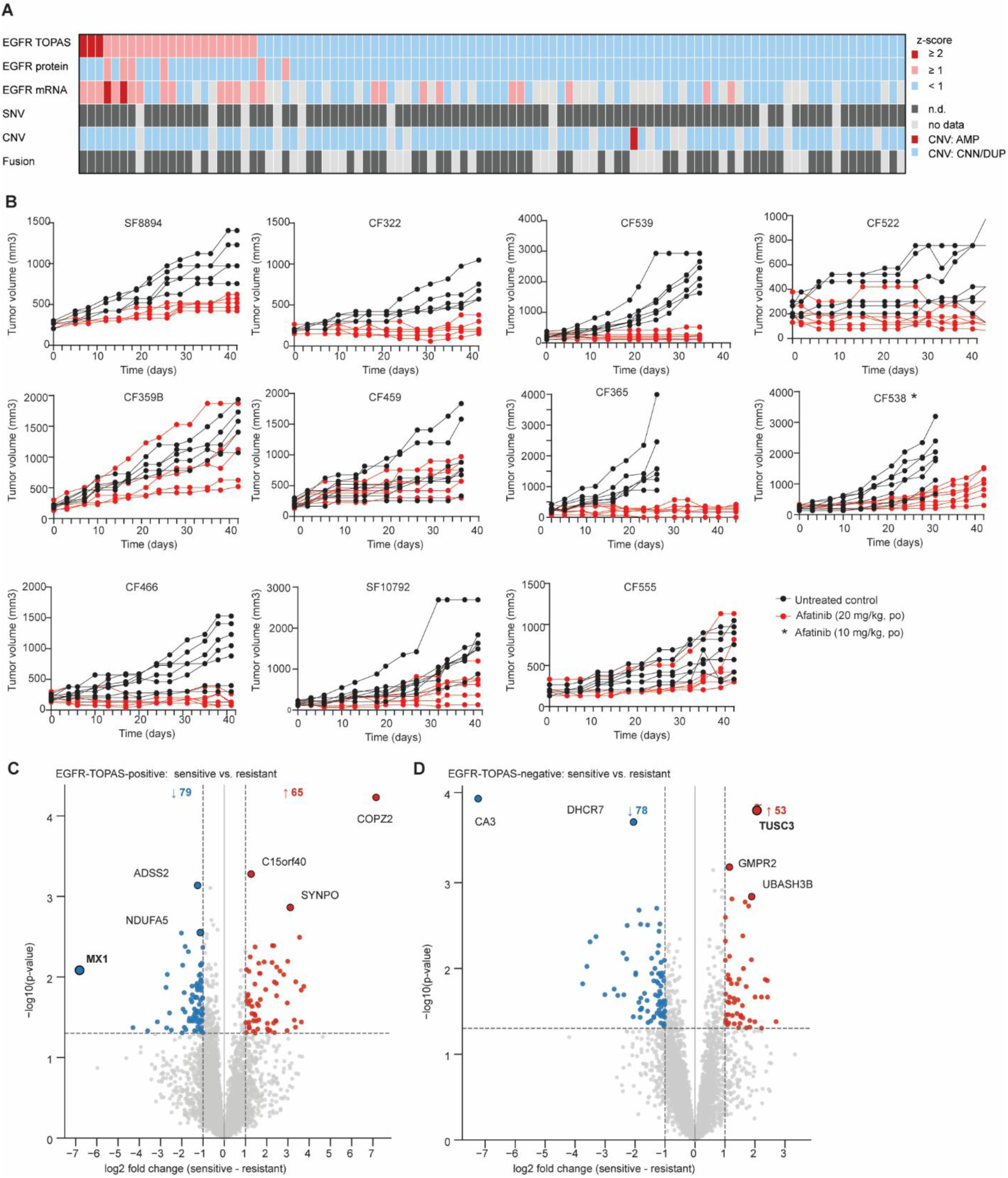
Stratification of chordoma patients for EGFR-targeted therapy. (A) Heatmap depicting EGFR pathway activity and genomic/transcriptomic alterations sorted by EGFR TOPAS kinase activity score from highest (left) to lowest (right) across all chordoma tumor specimen. Each column represents one patient (patient ID shown above). Rows show, from top to bottom: EGFR kinase activity score (TOPAS), EGFR protein abundance (z-score), EGFR mRNA expression (FPKM z-score), and somatic SNV, CNV, and gene fusion status. For quantitative rows, red indicates z-score ≥ 2, light red indicates z-score ≥ 1, light blue indicates z-score < 1, and grey indicates missing data. For genomic alteration rows, red indicates amplification (AMP); light blue indicates duplication (DUP), copy-number neutral (CNN), or deletion (DEL); dark grey indicates not determined (n.d.); light grey indicates not available (NA). (B) Tumor growth (volume) curves of PDX models treated with afatinib (20 mg/kg or 10 mg/kg daily, oral) in red or untreated in black for 42 days (or until the control group reached a mean tumor volume of 1,500 mm3), n=5-7 animals per group. (C,D) Volcano plot of differential expressed proteins in the EGFR-positive (C) or EGFR-negative (D) sensitive vs. resistant chordoma models (cell lines and PDX).

